# Promoterless Transposon Mutagenesis Drives Solid Cancers via Tumor Suppressor Inactivation

**DOI:** 10.1101/2020.08.17.254565

**Authors:** Aziz Aiderus, Ana M. Contreras-Sandoval, Amanda L. Meshey, Justin Y. Newberg, Jerrold M. Ward, Deborah Swing, Neal G. Copeland, Nancy A. Jenkins, Karen M. Mann, Michael B. Mann

## Abstract

A central challenge in cancer genomics is the systematic identification of single and cooperating tumor suppressor genes driving cellular transformation and tumor progression in the absence of oncogenic driver mutation(s). Multiple *in vitro* and *in vivo* gene inactivation screens have enhanced our understanding of the tumor suppressor gene landscape in various cancers. However, these studies are limited to single or combination gene effects, specific organs, or require sensitizing mutations. In this study, we developed and utilized a *Sleeping Beauty* transposon mutagenesis system that functions only as a gene trap to exclusively inactivate tumor suppressor genes. Using whole body transposon mobilization in wild type mice, we observed that cumulative gene inactivation can drive tumorigenesis of solid cancers. We provide a quantitative landscape of the tumor suppressor genes inactivated in these cancers, and show that despite the absence of oncogenic drivers, these genes converge on key biological pathways and processes associated with cancer hallmarks.

## Introduction

Recent genomic analysis of physiologically and histologically normal tissues such as eyelid epidermis and esophageal squamous epithelia show that these tissues tolerate relatively high levels of mutations, typically within known tumor suppressor genes (Martincorena et al., 2018; Martincorena et al., 2015; Yokoyama et al., 2019). *Sleeping Beauty* (SB) insertional mutagenesis (Ivics et al., 1997) is a powerful forward genetic tool used to perform genome-wide forward genetic screens in laboratory mice for cancer gene discovery (Collier and Largaespada, 2007; Copeland and Jenkins, 2010; Dupuy et al., 2005; Dupuy et al., 2009; Mann et al., 2014a; Mann et al., 2016b; Mann et al., 2012; Mann et al., 2015; Mann et al., 2014b; Rangel et al., 2016; Takeda et al., 2015). We recently conducted and reported an SB screen to model the development of cutaneous squamous cell carcinoma *in vivo*, and noted that approximately 30% of tumors formed without any oncogenic transposon insertions, albeit with extended latency. This finding raises two questions: (i) can exclusive loss of tumor suppressor genes *per se* drive tumorigenesis? and (ii) how do the kinetics of tumorigenesis in this context compare to tumors that develop via alterations in both tumor suppressor and oncogenes?

In this study, we generated a series of new SB transposon alleles to explore the hypothesis that cumulative loss of tumor suppressor genes is sufficient to drive the initiation and progression of systemic tumorigenesis. To this end, we modified key features of an existing high copy pT2/Onc2 transposon allele (Dupuy et al., 2005) and engineered a new transposon construct that functions as a gene trap to inactivate gene expression. Whole body transposition of this modified high-copy, gene-trap transposon (pT2/Onc2.3 or SB-Onc2.3) via inducibly expressed SB transposase was sufficient to initiate and progress a variety of tumor types *in vivo*. Here, we prioritized solid tumors of the skin, lung and liver and employed high-throughput SBCapSeq approaches (Mann et al., 2016b; Mann et al., 2015) to identify genome-wide SB mutations and define recurrently mutated, statistically significant candidate cancer drivers (CCDs) from bulk tumors containing more SB mutations expected by chance using the SB Driver Analysis (Newberg et al., 2018a) statistical framework. We provide a quantitative genetic landscape of the transposon insertions in these tissues and demonstrate that Ras signaling and members of the ubiquitin ligase complex are frequently inactivated, suggesting key roles of these biological process and pathways during the initiation and progression of solid cancers in the absence of selected oncogenic events.

## Results

### Harnessing an SB gene-trap allele for forward genetic screens

#### Whole-body mutagenesis using high-copy, gene-trap only SB transposons in wild type mice

We derived a novel gene-trap-only SB transposon allele, pT2/Onc2.3 (a.k.a. pT2/SB-GT-MBM102, **Figure 1a**), with two alterations to the pT2/Onc2 (Dupuy et al., 2005) transposon vector. First, the MSCV 5’LTR promoter and splice donor sequences were removed to disable the ability of the transposon to activate gene expression upon integration into the promoter or intron regions of a gene. Second, an additional bi-directional SV40-polyA signal sequence was added to enhance the transcriptional termination ability of the transposon upon integration into genes. These changes shortened the Onc2.3 SB transposon cargo to ∼1.6 kb (**Figure 1a**), which matches the natural size of the original fish SB(Izsvak and Ivics, 2004) and optimum size to maximize transposition in mammalian cells(Izsvak et al., 2000). Collectively, these modifications result in an SB transposon that can only disrupt gene expression via inactivation, and facilitates the identification of inactivated tumor suppressor genes that drive tumor initiation and progression *in vivo*.

**Figure 1:**
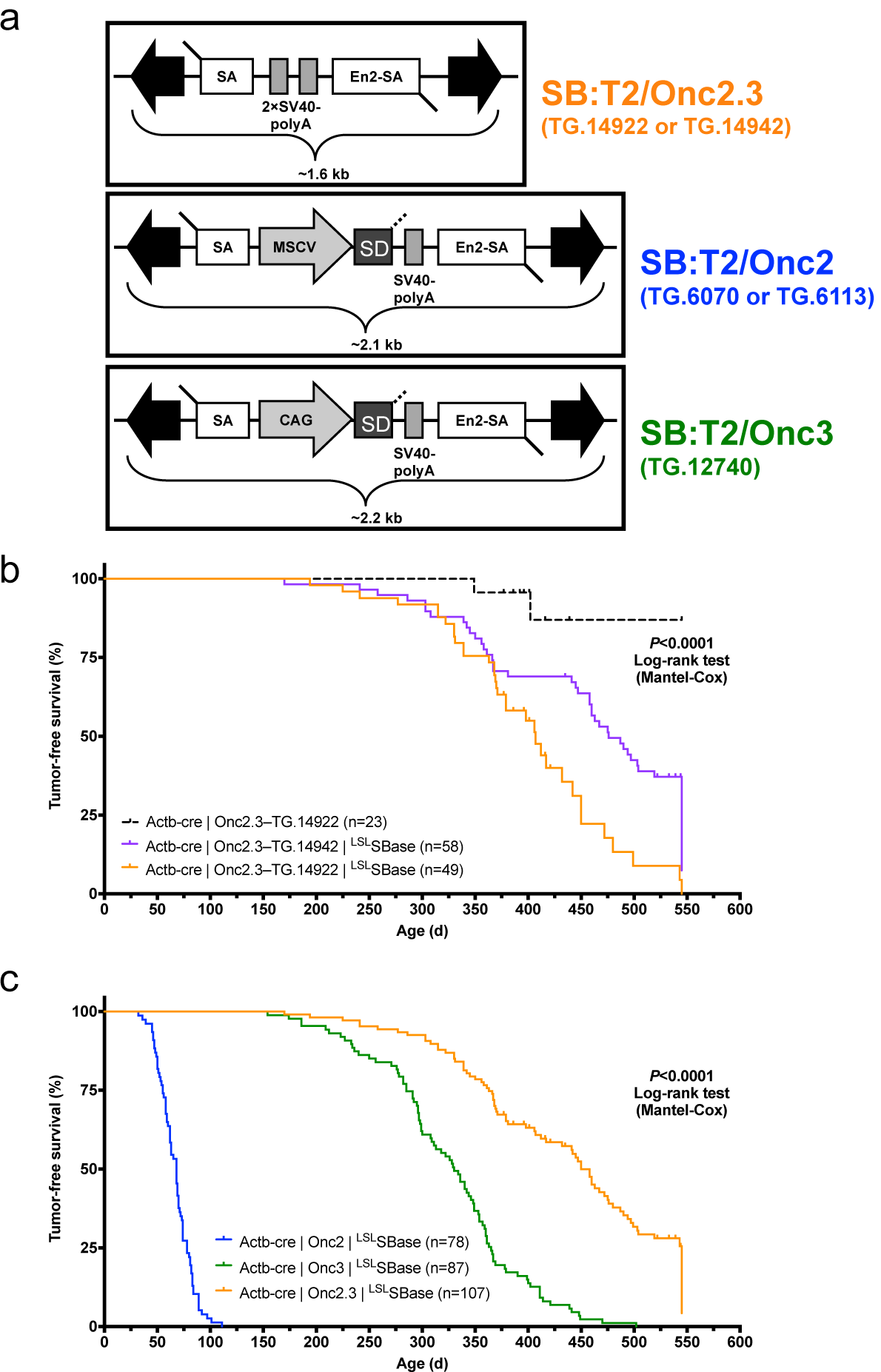
Whole-body, gene-trap exclusive SB transposon mutagenesis results in tumor formation in SB-Onc2.3 mice. (**a**) Comparing *Sleeping Beauty* transposons used in this study (T2/SB-GT, T2/Onc3) or in a similar companion study (T2/Onc2). Abbreviated allele designations for founder lines used in this work are shown to the right of each vector. (**b**) Kaplan-Meier survival plots comparing experimental and control SB-Onc2.3 cohorts (Mantel-Cox log-rank test, *P* < 0.0001, **Supplementary Figure 1**). Mice in all SB-Onc2.3 cohorts with active SB mobilization developed solid tumors at multiple organ sites, however SB-Onc2.3-TG.14922 mice (orange line) had significantly decreased survival compared to the SB-Onc2.3-TG.14942 mice (purple line; Mantel-Cox log-rank test, *P* = 0.0015). (**c**) Kaplan-Meier survival plots comparing the relative tumor latency between various SB cohorts with whole-body mobilization by a conditionally activated from the *Rosa26*-^LSL^SBase allele to the *Rosa26*-1lox-SBase allele by whole body Beta-actin-driven *Cre* expression, including SB-Onc2 mice (combined data from TG.6070 and TG.6113 alleles containing an MSCV-promoter (Mann et al., 2016b)), SB-Onc3 (data from TG.12740 allele containing a CAG-promoter (Aiderus et al., 2019)), and SB-Onc2.3 (combined data from TG.14922 and TG.14922 alleles containing no promoter) (Mantel-Cox log-rank test, *P* < 0.0001, **Table 1**).

After pT2/Onc2.3 plasmid linearization (**Supplementary Figure 1a**) and microinjection into pronuclei, 56 of the resulting 101 live-born and weaned progeny were screened by Southern blot and found to carry at least one copy of the transposon transgene (**Supplementary Figure 1b**). We reasoned that mice with the highest transgene copy number possible would allow for a maximal number of independent integration events per cell, a strategy that facilitates identification of cooperating TSGs in malignant transformation. Initially, five transgenic lines were selected to test for germ-line transmission by backcross breeding to C57BL/6J mice (**Supplementary Figure 1b**). All five lines transmitted to offspring, but only three T2/Onc2.3 lines, TG.14913, TG.14922, and TG.14942, segregated a transposon concatemer as a discreet locus, suggesting they arose from a single event leading to high-copy concatemer integration into the respective donor genomes.

**Table 1:** SB allele-specific tumor spectrum by whole-body, transposon-mediated mutagenesis in wild type mice. Distribution of cancer types between promoter-driven (SB-Onc3, SB-Onc2) and promoter-less (SB-Onc2.3) transposons. Complete Table 1 source data may be downloaded from Figshare http://dx.doi.org/10.6084/m9.figshare.12811685.

| SB Cohort<br>Promoter<br>Reference | SB-Onc2.3<br>no promoter<br>This study |  | SB-Onc3<br>CAG promoter<br>Aiderus et al., 2019 |  | SB-Onc2<br>MSCV promoter<br>Mann et al., 2016 |  |
| --- | --- | --- | --- | --- | --- | --- |
| Tumor classification | tumors | mice | tumors | mice | tumors | mice |
| Skin Squamous Cell Carcinoma | 5 | 4 (3%) | 52 | 20 (24%) | 0 | 0 (0%) |
| Skin Keratoacanthoma | 1 | 1 (1%) | 9 | 8 (10%) | 0 | 0 (0%) |
| Hepatocellular Carcinoma | 2 | 2 (2%) | 9 | 9 (11%) | 0 | 0 (0%) |
| Hepatocellular Adenoma | 4 | 3 (2%) | 79 | 42 (50%) | 0 | 0 (0%) |
| Adenoma, Multiple | 25 | 16 (12%) | 39 | 31 (37%) | 0 | 0 (0%) |
| Adenocarcinoma, Multiple | 3 | 3 (2%) | 9 | 8 (10%) | 0 | 0 (0%) |
| Carcinoma, Multiple | 0 | 0 (0%) | 6 | 6 (7%) | 0 | 0 (0%) |
| Sarcoma | 0 | 0 (0%) | 6 | 6 (7%) | 0 | 0 (0%) |
| Papilloma | 1 | 1 (1%) | 4 | 3 (4%) | 0 | 0 (0%) |
| Astrocytoma | 34 | 34 (26%) | 3 | 3 (4%) | 0 | 0 (0%) |
| Hemangiosarcoma | 0 | 0 (0%) | 3 | 3 (4%) | 0 | 0 (0%) |
| Hemangioma | 1 | 1 (1%) | 2 | 2 (2%) | 0 | 0 (0%) |
| Fibrosarcoma | 0 | 0 (0%) | 2 | 2 (2%) | 0 | 0 (0%) |
| Metastasis | 0 | 0 (0%) | 2 | 2 (2%) | 0 | 0 (0%) |
| Mast Cell Tumor | 0 | 0 (0%) | 1 | 1 (1%) | 0 | 0 (0%) |
| Trichoepithelioma | 0 | 0 (0%) | 1 | 1 (1%) | 0 | 0 (0%) |
| Schwannoma | 1 | 1 (2%) | 0 | 0 (0%) | 0 | 0 (0%) |
| Hepatoblastoma | 1 | 1 (1%) | 0 | 0 (0%) | 0 | 0 (0%) |
| Medulloblastoma | 0 | 0 (0%) | 0 | 0 (0%) | 6 | 6 (8%) |
| Lymphoma | 1 | 1 (2%) | 2 | 2 (2%) | 28 | 28 (36%) |
| Leukemia | 0 | 0 (0%) | 0 | 0 (0%) | 78 | 78 (100%) |
| Histocytic Sarcoma | 0 | 0 (0%) | 8 | 8 (10%) | 0 | 0 (0%) |
| **Totals** | **79** | **107** | **237** | **87** | **112** | **78** |
| **Average tumors per mouse** | **0.74** |  | **2.72** |  | **1.44** |  |
| **Average tumor latency (days)** | **450** |  | **330** |  | **70** |  |

The three SB-Onc2.3 transgenic lines contain approximately the same number of copies as other T2/Onc2 high-copy lines, with estimated copy number of ∼400 per transgene array (**Supplementary Figures 2a-b, 3a-b**), and each line could be maintained as transgenic homozygotes without loss of viability, fertility, or other obvious phenotypes. However, similar to what was observed with high-copy pT2/Onc2 carrier mice(Dupuy et al., 2005), when combined with a constitutively active SBase allele all three T2/Onc2.3 transgenic lines had loss of viability with statistically significant reduced numbers of live-born double heterozygous carriers (**Supplementary Figure 4**). To circumvent lethality in TG.14922 and TG.14942 lines, we used a conditional ^LSL^ SBase and *Actb*-Cre allele strategy (Aiderus et al., 2019; Mann et al., 2016b) that fully restored viability in the resulting triple heterozygous carrier mice (**Supplementary Figure 4a-b**). When combined with conditional ^LSL^ SBase and *Actb*-Cre alleles, T2/Onc2.3 transgenic carrier mice (SB-Onc2.3 hereafter), exhibited high rates of whole-body transposition that resulted in tumor formation (**Figure 1b** and **Supplementary Figure 5a-c**).

#### Systemic, whole body Onc2.3 transposition extends latency of tumor development

We generated 107 SB-Onc2.3 (n=58 Actb-Cre|Onc2.3-TG.14942; n=49 Actb-Cre|Onc2.3-TG.14922) and 23 control (Actb-Cre|Onc2.3-TG.14922) mice and aged them for a maximum of 550 days (1.5 years). At the predetermined endpoint, while only two control mice developed masses, many of the Onc2.3 developed tumors (**Figure 1b**). Tumor-free survival of both SB-Onc2.3 cohorts was significantly reduced compared to control cohorts (Kaplan-Meier log-rank test, *P*<0.0001, **Figure 1b**). Tumor latency was significantly reduced in the TG.14922 cohort compared with the TG.14942 cohort (Kaplan-Meier log-rank test, *P*<0.0001, **Figure 1b**) despite having indistinguishable tumor spectrums. Some of the mice censored in the survival analysis were observed to have various cancers when necropsied and/or histologically evaluated, suggesting our data represent a lower limit to the full tumor spectrum in SB-Onc2.3 mice.

The combined SB-Onc2.3 cohort exhibited a significantly longer mean tumor latency (Log rank test, *P*<0.0001; **Figure 1c**) compared to similarly bred wild type SB-Onc2 (Mann et al., 2016b) or SB-Onc3 (Aiderus et al., 2019) cohorts. The tumor latency and spectrum of SB-Onc2.3 mice were more similar to SB-Onc3 (Aiderus et al., 2019) mice, with few hematopoietic tumors and a similar wide variety of solid tumor types, compared with SB-Onc2, including skin squamous cell carcinomas (cuSCC)/keratoacanthoma (cuKA) and hepatocellular adenomas (HCA)/carcinomas (HCC) (**Table 1** and **Supplementary Table 1**). Unexpectedly, SB-Onc2.3 mice exhibited a significantly larger proportion of early-stage, low-grade solid tumor types, including spinal cord and cerebellar astrocytomas (ACT) and lung alveolar adenoma (LUAA) and adenocarcinoma (LUAC) (**Supplementary Figure 6** and **Supplementary Table 1**). However, with respect to transposon copy number, the average number of SB insertions observed per tumor genome in SB-Onc2.3 mice was more similar to SB-Onc2 compared with low-copy SB-Onc3 (**Supplementary Figure 3-4**), suggesting that copy-number *per se* does not explain the differences in latency and tumor spectrum.

To better understand the genetic events driving these SB-Onc2.3 solid tumors, we generated genomic libraries from a wide variety of SB-Onc2.3-induced solid cancers and used high-throughput SBCapSeq (Mann et al., 2016b; Mann et al., 2015) to identify genome-wide SB insertions. Similar to previous reported results in SB-Onc2(Mann et al., 2016b) and SB-Onc3(Aiderus et al., 2019), the SB-Onc2.3 libraries were highly reproducible (**Supplementary Figure 7** and **Supplementary Table 2**).

### Solid tumors driven by TSG inactivation in SB-Onc2.3 mice

#### Landscape of cuSCC driven by transposon-mediated tumor suppressor inactivation

Skin tumors comprised 4% of the total number of tumors collected and histologically verified in SB-Onc2.3 mice, suggesting that inactivation of tumor suppressor genes is sufficient to drive keratinocyte initiation and progression to frank cuSCC *in vivo*. We recently reported in an SB-driven cuSCC model that activating SB insertions into *Zmiz1, Zmiz2*, or *Mamld1* oncoproteins are collectively observed in over two-thirds of cuSCC tumors sequenced using SBCapSeq (Aiderus et al., 2019). This suggests that the initiation and progression of approximately one-third of skin tumors *in vivo* do not require positive selection for proto-oncogenes. We hypothesized that tumors without activating oncogenic insertions in the *Zmiz1 or Zmiz2* genes would have higher frequencies of inactivation of the tumor suppressor genes found in the Onc2.3 system. We used SBCapSeq to quantitatively define the gene insertions in 6 SB-Onc2.3 skin tumors with confirmed cuSCC diagnosis (**Supplementary Tables 3-4**) (Mann et al., 2016b). As expected, no activating SB insertions greater than background rates (≥1 SBCapSeq read) were observed in the proto-oncogenes *Zmiz1, Zmiz2*, or *Mamld1* previously reported in our SB-Onc3-driven cuSCC or cuKA cohorts(Aiderus et al., 2019). In contrast, the SB-Onc2.3 cuSCC tumors had recurrent, high read-depth inactivating insertions in the *Rasa1, Trip12, Cul3, Cux1, Tcf12, Nf1, Kmt2c, Ncoa2*, and *Crebbp* genes (**Figure 2**). Pathway analysis using Enrichr software (Chen et al., 2013; Kuleshov et al., 2016) and KEGG database revealed enrichment of selected SB insertions in genes associated with MAPK signaling pathway (*Abl2, Nf1, Rasa1, Stk4*; *P*=0.003) and Ras signaling pathways (*Nf1, Rasa1, Stk4, Tgfb2*; *P*=0.008). Based on these data, we revisited our previously reported sequencing data from the SB|Onc3 cuSCC tissues (Aiderus et al., 2019). Indeed, we observed that Zmiz1/2-negative cuSCC tumors had dramatically higher inactivating SB insertions in *Trip12, Kmt2c, Rasa1, Cul3* and *Arid1b* in ZMIZ1/2-negative cuSCC tumors relative to ZMIZ1/2-positive cuSCC genomes (**Figure 2**). Notably, almost all (95%, 22/23) of the Zmiz1/2-negative cuSCC contained single or cooperating mutations in those five genes (**Figure 2**), compared to 63% (27/43) of the ZMIZ1/2-positive cuSCC genomes. While the frequencies of individual gene hits within the Zmiz1/2-negative cuSCC genomes derived from either Onc3 or Onc2.3 SB transposon lines varied, similar cooperating relationships among the putative TSGs were conserved (**Figure 2**). We conclude that the Onc2.3 transposon-derived skin tumors develop via cooperating inactivation of tumor suppressor genes. This supports our prior observation that approximately one-third of SB-driven skin masses are not driven by activating SB insertions in proto-oncogenes (Aiderus et al., 2019).

**Figure 2:**
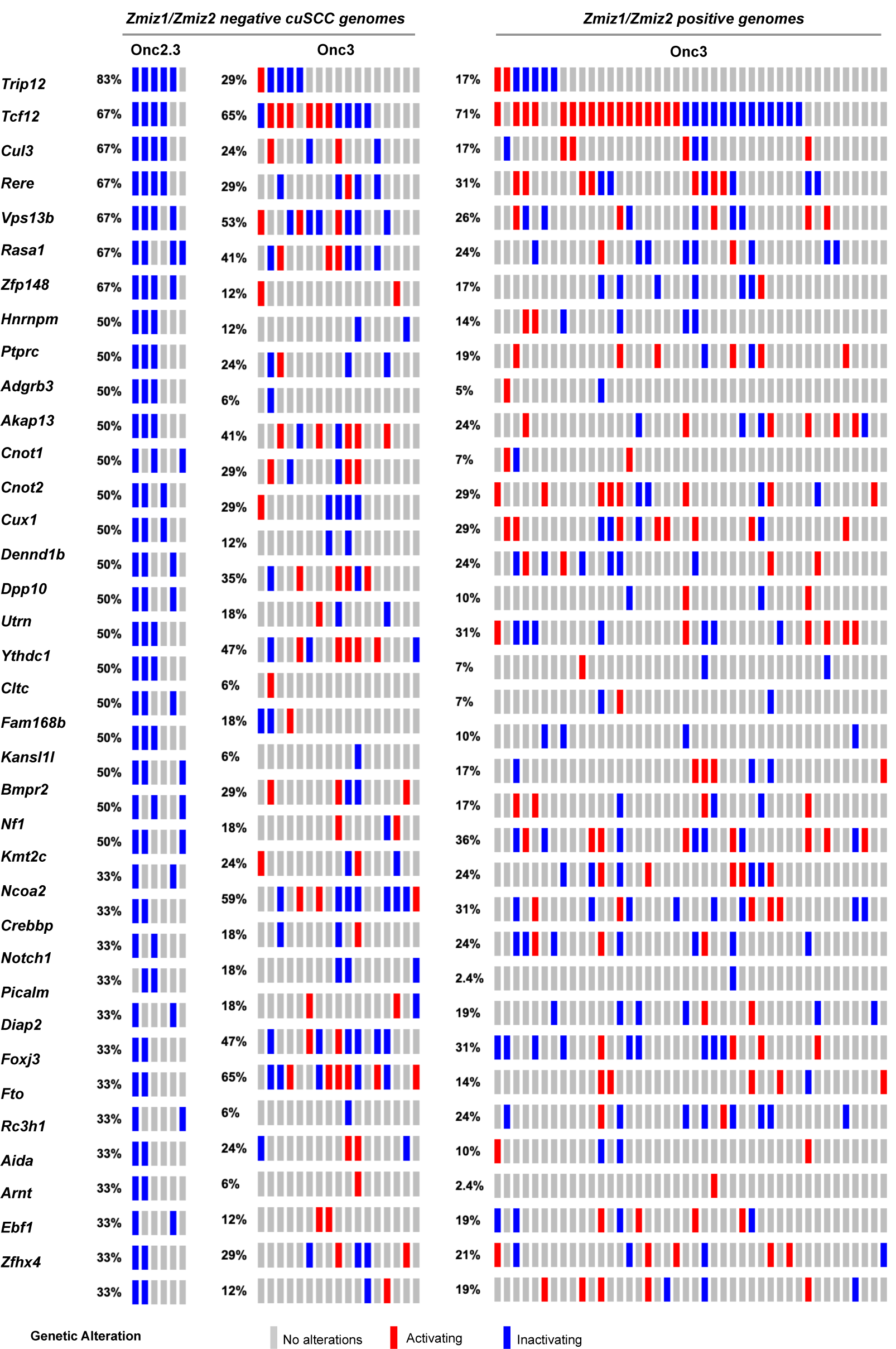
SB-driven cuSCC by cumulative tumor suppression. (**a**) Recurrent inactivating SB insertions in Onc2.3-induced cuSCC genomes (n=6). (**b**) Comparative waterfall plots of trunk driver genes from *Zmiz1*/*Zmzi2*-independent SB-Onc2.3 and SB-Onc3 (n=23; left panel) and *Zmiz1*/*Zmzi2*-dependent SB-Onc3 (n=43; right panel) cuSCC tumors. (**c**) *Zmiz1*/*Zmiz2*-independent tumor segregated by SB cohort, SB-Onc3 (n=17; left panel) and SB-Onc2.3 (n=6; right panel).

#### Landscape of lung cancer driven by transposon-mediated tumor suppressor inactivation

Approximately one-third of lung adenocarcinomas contain oncogenic mutation in KRAS (Suh et al., 2016), and lung tumorigenesis is usually modeled in mice using oncogenic mutant *Kras* alleles as initiating events (Haigis, 2017). Using SB mutagenesis, we successfully modeled lung alveolar adenoma (LUAA) and adenocarcinoma (LUAC), demonstrating that oncogenic mutant *Kras*, or other strong oncogenic drivers, is not required for lung tumor initiation in the mouse. To dissect the selected SB events driving early lung LUAA/LUAC tumorigenesis, we applied the SBCapSeq methodology to analyze the insertion profiles of tumors driven by the transposons Onc2.3 (**Supplementary Table 5**) or Onc3 (**Supplementary Table 6**; previously reported (Aiderus et al., 2019)), which can either inactivate alone or activate and/or inactivate gene expression, respectively, upon transposon insertion. In the Onc3 lung LUAA/LUAC tumors, 70% had directional, activating insertions in *Rasgrf1*, encoding the guanine nucleotide exchange factor RASGRF1 which functions to activate Ras signaling by exchanging GDP for GTP (**Figure 3, Supplementary Figure 8a-b, and Supplementary Table 6**). Results from 454 sequencing confirmed activating *Rasgrf1* insertions among the 29 Onc3 LUAA/LUAC tumors, including 10 large tumors from flash frozen genomic DNA which were also sequenced by SBCapSeq, and 19 additional smaller tumors isolated from FFPE tumor tissues (**Supplementary Figure 9** and **Supplementary Tables 7-9**). In contrast, no insertions in *Rasgrf1* were observed in SB-Onc2.3 LUAA/LUAC tumors. Several genes including *Rbms3, Sik3, Cul3, Trip12, Rock1, Zfp292* and *Rasa1* were more commonly inactivated in Onc2.3 compared to Onc3-driven lung tumors. Pathway analysis using the Enrichr KEGG pathway analysis and gene ontology biological processes enrichment analysis revealed statistically significant enrichment of genes with SB insertions associated with focal adhesion pathway (*Rock1, Rasgrf1, Pten*, and *Pik3r1*; *P*<0.0001), Ras signaling pathway (*Rock1, Rasgrf1*, and *Pik3r1*; *P*<0.0001) and regulation of protein ubiquitination (*Cul3* and *Trip12*; *P*=0.001). Notably, *Cul3* and *Trip12* — members of the E3 ubiquitin ligase complex — were exclusively inactivated in the Onc2.3 but not Onc3–driven lung tumors. Among the targets of TRIP12 is ASXL1, a tumor suppressor with roles in Polycomb-induced gene silencing. Two SNVs in *Asxl1* were identified in mouse lung adenomas induced by MNU in the background of a Kras^G12D^ mutation (Westcott et al., 2015). Izumchenko *et al*. described mutations in *ASXL1* and other genes involved in DNA damage and chromatin remodeling identified in the lungs of human patients with atyptical adenomatous hyperplasia, an initiating event in adenocarcinoma development(Izumchenko et al., 2015). RBMS3, RNA Binding Motif Single Stranded Interacting Protein 3, is a validated TSG in human lung SCC where it acts primarily via regulation of c-Myc (Liang et al., 2015). CUL3 is a promiscuous E3 ubiquitin ligase that partners with multiple proteins to regulate protein turnover of many targets in a context specific manner. CUL3 plays an important role in ubiquitination of KEAP1, a negative regulator of the transcription factor NRF2 which enacts a transcriptional program important in protecting lung cancer and other tumor types from oxidative stress (Zhang et al., 2005). These data suggest that in the Onc3–driven lesions, activation of *Rasgrf1* contributes to rapid lung tumorigenesis, while in Onc2.3–driven lung tumorigenesis, the inactivation of cooperative tumor suppressor genes drives disease after long latency, potentially through modulating protein turnover and transcriptional reprogramming. Further studies are needed to show whether these mechanisms may represent functional redundancy with key pathways initiated by oncogenic *Kras* to promote lung cancer in mice.

**Figure 3:**
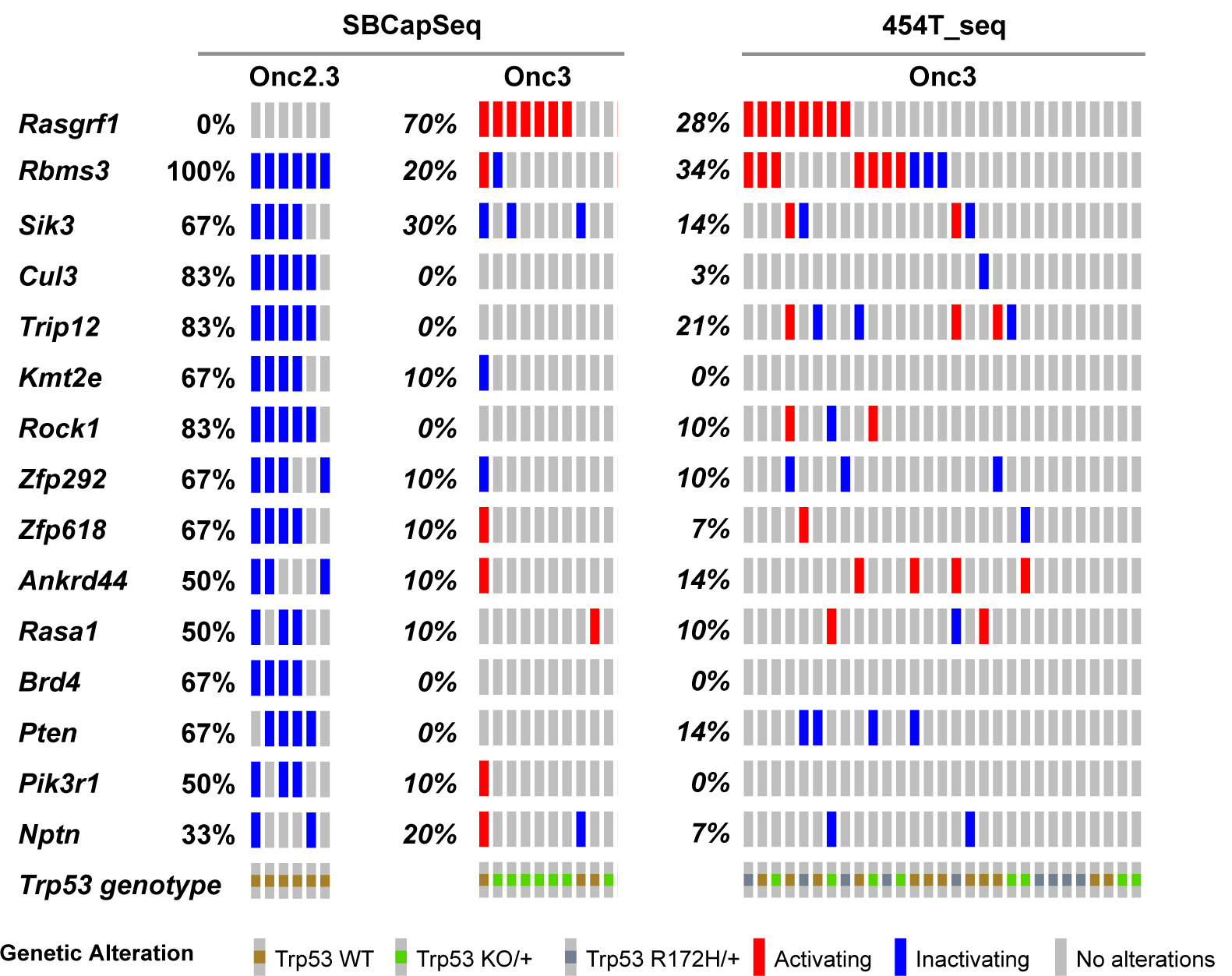
SB-driven lung cancer by cumulative tumor suppression. Recurrent inactivating SB insertions in Onc2.3 (n=6 using SBCapSeq) and Onc3 (n=10 using SBCapSeq; n=29 using 454T sequencing) induced LUAA genomes.

#### Landscape of liver cancer driven by transposon-mediated tumor suppressor inactivation

Previously we generated hepatocellular adenomas (HCAs) and adenocarcinomas (HCC) driven by the SB-Onc3 transposon in the presences of mutant or wild type *Trp53* alleles (Newberg et al., 2018a; Newberg et al., 2018b). From 95 histologically confirmed HCAs (HCA; **Supplementary Table 2**), we identified 43 trunk drivers, including proto-oncogenes *Hras, Kras, Rlt1*/*Rian* and tumor suppressor genes *Pten, Adk*, and *Zbtb20* (**Supplementary Tables 10–12**). Most of these genes were identified in *Trp53* mutant and wild type cohorts, suggesting that their role in driving HCA is *Trp53*-independent. Further, the majority of the putative driver genes identified in our HCA tissues were also reported in SB-driven hepatocellular carcinoma (HCC)(Bard-Chapeau et al., 2014; Dupuy et al., 2009; Keng et al., 2013; O’Donnell et al., 2012; Rogers et al., 2013), suggesting that many of the initiating genes enriched during earlier stages of hepatocyte transformation *in vivo* are maintained in HCC (**Supplementary Fig. 10**). SBCapSeq analysis of the 4 HCA and 2 HCC tumors from SB-Onc2.3 mice revealed no activating insertion patterns in the genomes from these tumors and the absence of hits in *Hras, Kras*, and Rlt1/Rian (**Supplementary Table 13**). However, we corroborated the tumor-suppressive roles of insertions in *Adk and Zbtb20*, and in *Nipbl, Pdlim5, Ppp1r12a, Tnrc6b, Brd4, Cul3, Ctnna3, Elavl1, Gphn, Nfia, Ptpn12, Taok3*, and *Rasa1* (**Figure 4**). Using Enrichr KEGG pathway analysis and gene ontology biological processes enrichment analysis revealed statistically significant enrichment of genes with SB insertions associated with ubiquitin mediated proteolysis (*Cul3, Cul5*, and *Trip12*; *P*<0.0001), growth factor signaling (*Ep300, Itgb1*, and *Rasa1*; *P*<0.001), scaffolding proteins involved in cell adhesion (*Dlg1, Itgb1*, and *Pdlim5*; *P*<0.001), and transcriptional regulators (*Brd4, Med13l, Nfia*, and *Nipbl*; *P*<0.0001). Notably, the inactivation of RASA1 suggests that the Onc2.3 transposon may activate Ras signaling via inactivation of the negative regulator of this pathway, in the absence of oncogenic Ras activation. Taken together, by using different transposon systems, we show that HCA in mice can be initiated by activation of oncogenes in combination with tumor suppressor inactivation, or exclusively by inactivation of cooperating tumor suppressors that converge on similar mechanisms co-opted to promote tumorigenesis.

**Figure 4:**
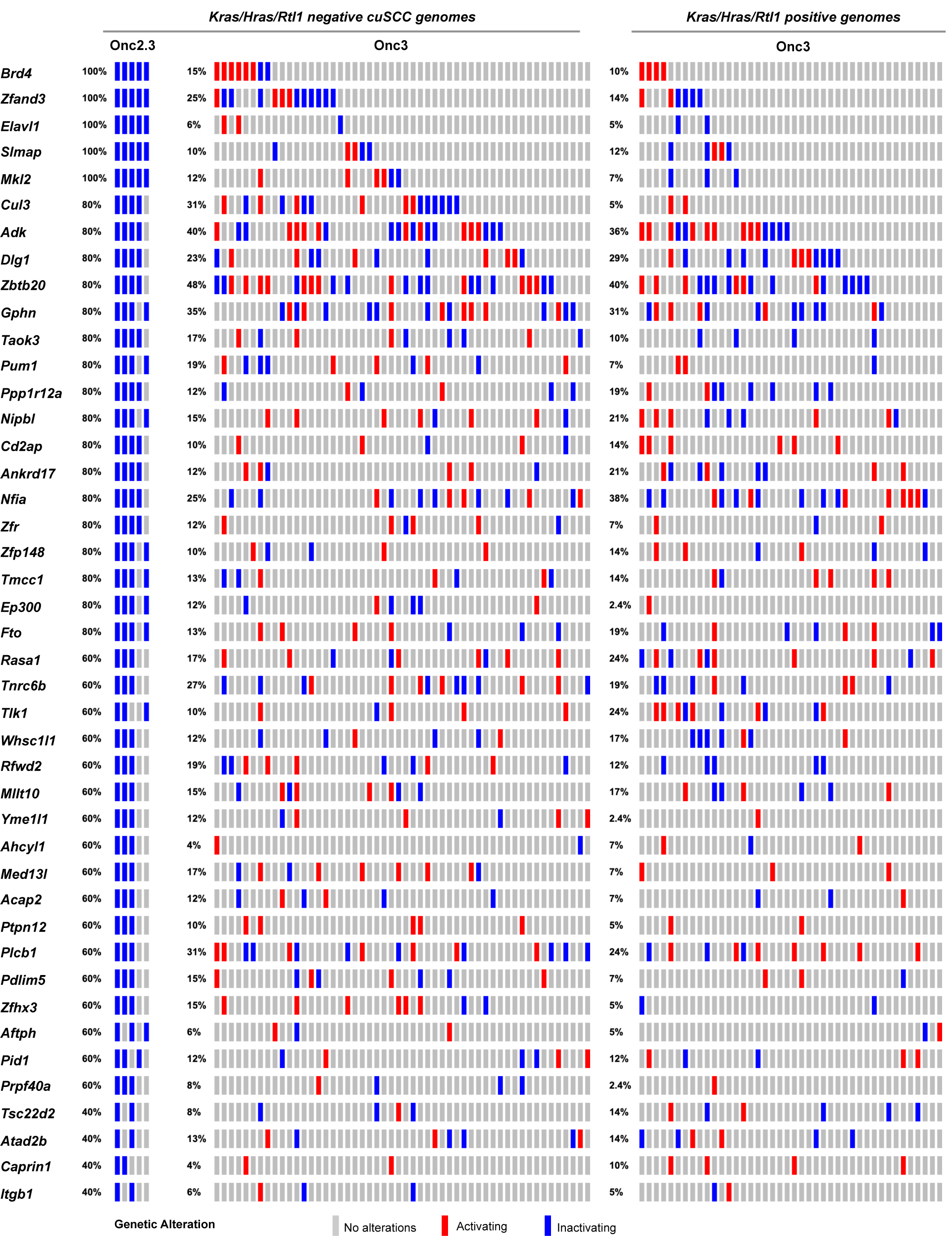
SB-driven liver cancer by cumulative tumor suppression. Recurrent inactivating SB insertions in Onc2.3 (n=6 using SBCapSeq) and Onc3 (n=52 *Kras*/*Hras*/*Rtl1* oncogene negative using 454T sequencing; n=42 *Kras*/*Hras*/*Rtl1* oncogene positive using 454T sequencing) induced HCA genomes.

**Figure 5:**
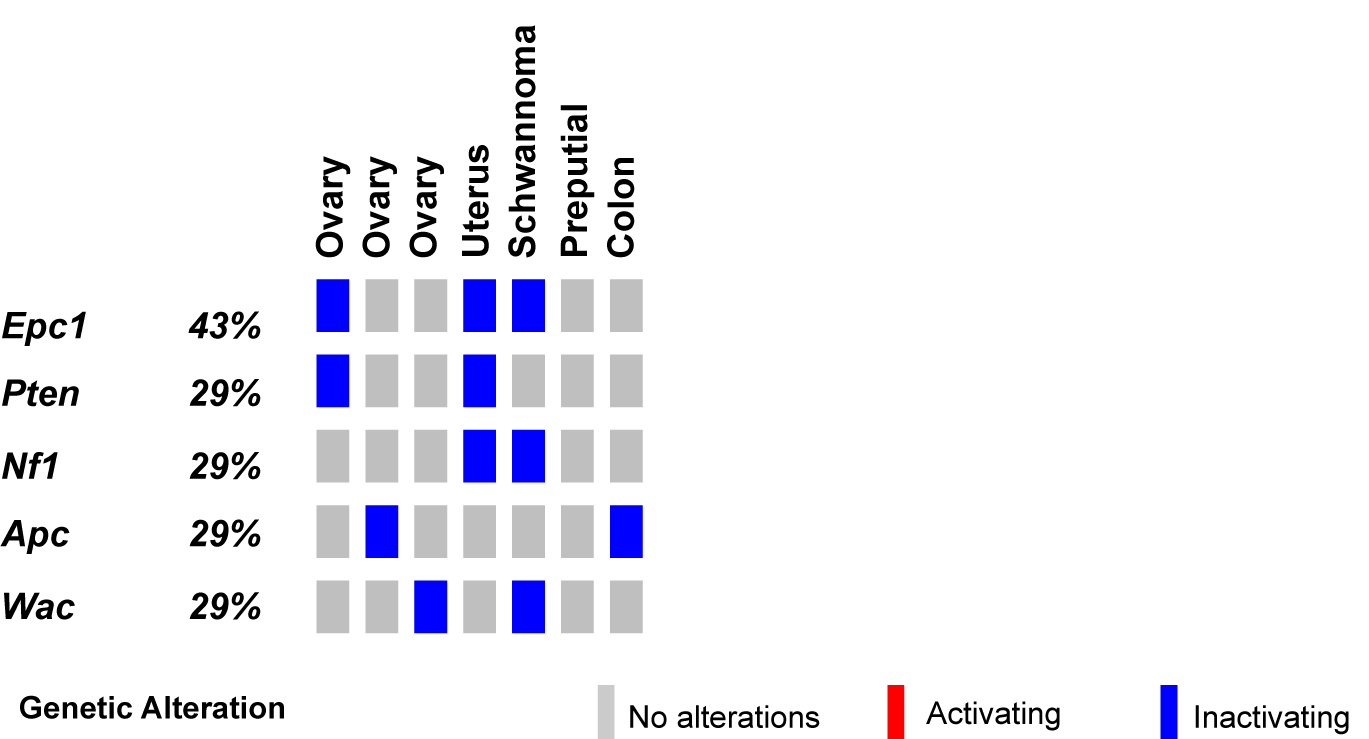
SB-driven gynecologic and other cancer histologies by cumulative tumor suppression. Recurrent inactivating SB insertions in Onc2.3-induced ovary, uterus, schwannoma, preputial gland, and colon tumors.

**Figure 6:**
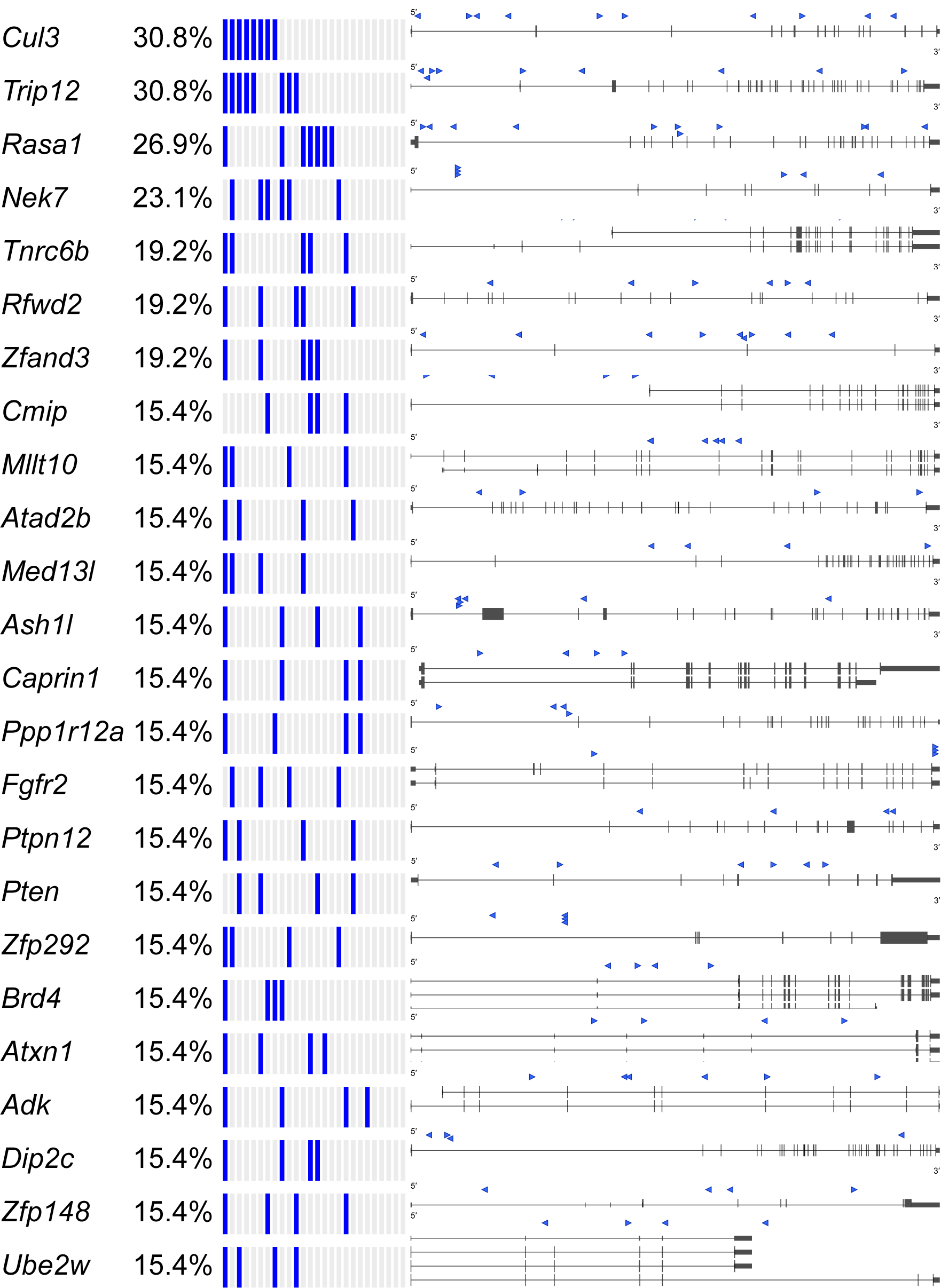
SB-Onc2.3-Pan-TSG driver discovery. Recurrent inactivating SB insertions into trunk drivers with >99 reads from 3 or more Onc2.3-induced cancer genomes (n=27). SB-Onc2.3-Pan-TSG driver gene SB insertion profiles are shown to the right. Literature References summarizing studies that provide in vivo evidence for validation of the TSGs identified in this study (**Supplementary Table 23**). Source image files may be downloaded from figshare http://dx.doi.org/10.6084/m9.figshare.12816365.

#### Landscape of other solid tumors driven by transposon-mediated tumor suppressor inactivation

Finally, we conducted SBCapSeq analysis on nine other SB-Onc2.3 mouse tumors from diverse tissues and histological classifications, including three ovarian tumors (with adenoma, papillary, and granulosa cell carcinoma subtypes), a uterine adenocarcinoma, a cerebellar astrocytoma, a colon polypoid adenoma, a preputial gland carcinoma, a renal tubular cell adenoma, and a Schwannoma of the cerebellum.

Individually, all but one specimen demonstrated reproducible (**Supplementary Table 3**) and clonal SB insertions sites, as evidenced by a range of low to very high read-depth-enriched TA-dinucleotide sites within the coding region of many known tumor suppressor genes such as *Apc, Brd4, Cul3, Cux1, Gnaq, Nf1, Pten, Rasa1, Rnf43*, and *Wac* (**Supplementary Table 14-15**). Notably, the astrocytoma sample presented no evidence of clonal selection, despite exhibiting the highest total number of SBCapSeq reads. This suggests that this mass may have other non-SB driven mechanisms contributing to tumor initiation (see **Supplementary Note 1**). Collectively, these results strongly suggest an SB-driven etiology, consistent with the studies in skin, lung, and liver described above, and in multiple previously reported studies (Aiderus et al., 2019; Mann et al., 2016b; Mann et al., 2012; Mann et al., 2015; Newberg et al., 2018a; Newberg et al., 2018b; Rangel et al., 2016; Takeda et al., 2015).

#### Common pathways selected in response to whole genome tumor suppressor gene inactivation

We observed *Brd4, Cul3*, and *Trip12* to be commonly mutated genes across different tumor types, suggesting that there may be common mechanisms perturbed by the Onc2.3 transposon to drive tumorigenesis in the absence of oncogenic events. Therefore, we extended our analysis to statistically define recurrent tumor suppressor genes across 27 SB-Onc2.3 tumors using SB capture sequencing (Mann et al., 2016b) and SB Driver analysis (Newberg et al., 2018a) (**Supplementary Table 16-18**). We reasoned that regardless of the tumor origin, a meta-analysis of selected SB insertion events enabled by the SB Driver Analysis (Newberg et al., 2018a) workflow may provide a quantitative means to explore the detection of potentially novel, cooperative TSGs. We identified 1,499 discovery-significant progression drivers, 476 genome-significant progression drivers (**Supplementary Table 17**), and 53 genome-significant trunk drivers (**Supplementary Table 18**) using the SB Driver Analysis statistical categorical framework (Newberg et al., 2018a) (see **Methods**). As expected, none of the sense-strand-oriented, recurrent SB insertions present in defined proto-oncogenes from tumors driven by SB transposons containing an internal promoter (see the SBCDDB (Newberg et al., 2018a; Newberg et al., 2018b)) were identified in any of the SB-Onc2.3 tumors. Further, meta-analysis using SB Driver Analysis, including all SB-Onc2.3 (n=27) and SB-Onc3 (n=10) lung tumors, showed that *Rasgrf1* was the only gene identified with an activating SB insertion pattern (**Supplementary Table 19**).

The Cancer Gene Census (COSMIC v91 (Bamford et al., 2004; Forbes et al., 2010; Sondka et al., 2018)) contains 313 genes with a designation of TSG and a unique mouse ortholog (**Supplementary Table 20**). The Onc2.3-Pan-TSG dataset contains 1,499 driver genes. We observed 78 genes in common between the CGC and the Onc2.3-Pan driver genes, which represent a statistically significant enrichment of shared genes than expected (**Supplementary Table 20**; Chi-square with Yates’ correction, χ^2^=104.742, two-tailed *P*<0.0001). This suggests that the genes identified in our screen are enriched for known TSGs that driver tumorigenesis. Selective and statistically significant enrichment of the Onc2.3-Pan-TSG dataset was also observed with the 1,004 TSGs listed in the Tumor Suppressor Gene Database (Zhao et al., 2016; Zhao et al., 2013) with a unique mouse orthologs. We observed 127 genes in common between the TSGdb-orthologs and the Onc2.3-Pan driver genes, which represent a statistically significant enrichment of shared genes than expected (**Supplementary Table 21**; Chi-square with Yates’ correction, χ^2^=32.662, two-tailed *P*<0.0001). Overall, 167 known TSGs were identified in the Onc2.3-Pan-TSG dataset, suggesting that the additional 1,333 remaining genes may represent potential novel TSG drivers (**Figure 7**), including *Trip12, Rbm33*, and *Zfp292*, with recurrent SB insertion sites in Onc2.3 tumors demonstrated insertion patterns predicted to inactivate target gene expression. Indeed, although not represented on either of the curated CGC or TSGdb lists, several genes (e.g., *Cep350* (Mann et al., 2015), *Ncoa2* (Aiderus et al., 2019; O’Donnell et al., 2012), and *Usp9x* (Perez-Mancera et al., 2012)) have all been observed in other SB tumor screens and experimentally validated to be TSGs within mouse models and/or human cancer cells.

**Figure 7:**
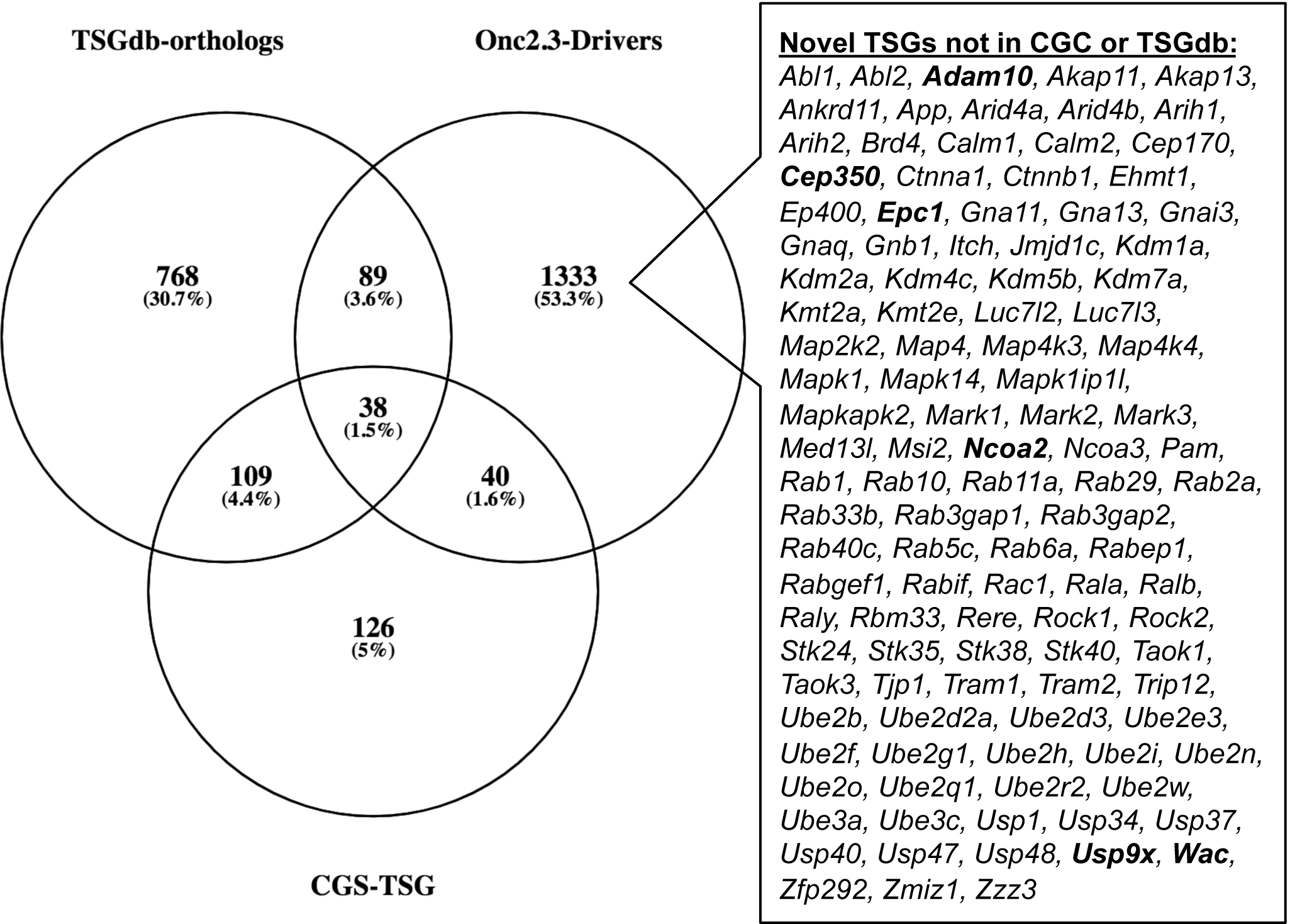
SB-Onc2.3-Pan-TSG driver enrichment with known TSGs datasets. The CGC TSG and TSGdb datasets overlap with 167 of the SB-Onc2.3-Pan-TSG driver genes. 111 putative novel TSGs not in the CGC TSG or TSGdb datasets, including validated TSGs (bold) described in the main text.

To gain further insights into novel pathways that lead to tumor initiation and/or progression via cooperating TSG only routes, we performed Enrichr biological pathway enrichment analysis (Chen et al., 2013; Kuleshov et al., 2016) using the 1,499 Onc2.3-Pan-TSG-Drivers. We observed overrepresentation of the Onc2.3-Pan-TSG-Drivers among the top 1% of all known biological processes with statistically significant enrichment in known pathways involved in cancer initiation and progression, including AR, EGF/ERGR, TGF-beta, PDGFB, mTOR, and VEGF signaling pathways, ubiquitin mediated proteolysis, chromatin modifying enzymes and organization, and RHO GTPase effectors (**Supplementary Table 22**). Similarly, Enrichr gene ontology enrichment analysis reaffirmed our manual curation, namely that genes with roles in diverse processes, including protein ubiquitination, cadherin binding, focal adhesion, and chromatin, are over-represented among the top 1% of all known biological processes (**Supplementary Table 22**). We conclude that this set of known and suspected TSGs represents a rich resource for prioritizing future efforts to catalogue the genes in the cancer genome that can drive tumor initiation and/or progression without oncogenic mutations.

## Discussion

In this study, we investigated whether *in vivo* tumor initiation and progression was possible solely by inactivation of tumor suppressor genes. We generated a new SB transposon allele that lacks the internal promoter present in all SB transposons used to-date in mouse models of cancer(Collier et al., 2005; Copeland and Jenkins, 2010; Dupuy et al., 2005; Dupuy et al., 2009; Newberg et al., 2018a; Newberg et al., 2018b). Here, we demonstrate that the Onc2.3 transposon restricts SB mutagenesis to act as a gene trap, selectively inactivating genes. While SB genetic screening studies have reported tumor suppressor genes in mouse models of cancer, these screens have often been conducted using strong oncogenic initiating mutations, including Kras^G12D^ (Mann et al., 2012; Perez-Mancera et al., 2012), Braf^V600E^ (Mann et al., 2015; Perna et al., 2015) and Apc loss (Starr et al., 2009; Takeda et al., 2015) or HBV expression (Bard-Chapeau et al., 2014). Importantly, we demonstrate that our Onc2.3 transposon system enables the evaluation of tumorigenesis *in vivo* by exclusive loss of tumor suppressor genes without introduction of an initiating oncogene (Katigbak et al., 2016; Zender et al., 2008). Notably, whole-body mutagenesis using the Onc2.3 transposon resulted in distinct cell types that were more susceptible to tumorigenesis than others. This may be due to the fact that exclusive tumor suppressor gene inactivation leads to a longer tumor latency in some tissues, or that some cell types in the mouse have robust mechanisms to withstand tumorigenesis without an exogenous or induced oncogenic hit. This latter phenomenon has been well-documented in the literature, particularly for mouse models of human cancers driven by oncogenic sensitizing events, including melanoma, pancreatic and colon cancers (Mann et al., 2012; Mann et al., 2015; Newberg et al., 2018a; Newberg et al., 2018b; Takeda et al., 2015), notably absent from the observed tumor types with the Onc2.3 promoterless transposon.

Mutagenesis methods to study skin tumorigenesis typically rely on inducing point mutations that target both tumor suppressors and oncogenes (Abel et al., 2009; Chitsazzadeh et al., 2016; Molina-Sanchez and Lujambio, 2019) or insertional mutations without point mutations (Mann et al., 2016b; Mann et al., 2015). In cuSCC tumors driven by the SB-Onc2.3 transposon, we noted a high frequency of gene inactivation in chromatin remodelers. This suggests that in the absence of an oncogenic hit, inactivation of chromatin remodeler genes can drive cuSCCs, presumably due to their dynamic ability to regulate expression of genes, miRNAs and lncRNAs involved in different biological processes and pathways intimately involved in tumorigenesis. Indeed, inactivation of chromatin remodeler genes has been shown to drive the initiation and progression of many different cancers (Andricovich et al., 2018; Cho et al., 2018; Dhar et al., 2018; Jia et al., 2018; Mathur et al., 2017; Wu et al., 2018). We recently reported functional evidence that shRNA-mediated knockdown of *KMT2C* or *CREBBP* in immortalized primary human keratinocytes was sufficient for cell transformation *in vitro* but was insufficient to promote tumor growth *in vivo*, suggesting that cuSCC development requires additional cooperating oncogenic events(Aiderus et al.). This is supported by the recent findings of Ichesi *et al*. who reported that heterozygous loss of *Crebbp* alone, or together with heterozygous loss of its paralog *Ep300*, cooperates with mutant Ras to drive keratinocyte hyper-proliferation *in vivo* (Ichise et al.). Furthermore, a recent genetic screen demonstrated that inactivation of *Fbxw7* or *Slc9a3* could transform mouse embryonic fibroblasts cells *in vitro* within the context of oncogenic *Kras*^G12D^, but was not sufficient to form tumors *in vivo* (Huang et al.). Our SB-Onc2.3 is also of relevance to human cuSCC, as the majority of patients are older individuals, and recent studies indicate that cancer driver mutations are tolerated in physiologically normal squamous tissues, even in relatively younger individuals (Martincorena et al., 2018; Martincorena et al., 2015). This is consistent with the observation in both this study (using SB-Onc2.3) and our previous study (using SB-Onc3) (Aiderus et al., 2019) that cumulative tumor suppressor inactivation over time may be an important and under recognized route to keratinocyte transformation initiation and cuSCC progression.

In both cuSCC and liver tumors, inactivation of the *Rasa1* gene, encoding a RAS-inhibiting GTPase, was a frequent, recurrent event. *Rasa1* inactivation was observed in a previous SB study investigating resistance mechanisms to FGFR inhibition in invasive lobular breast cancer (Kas et al.). Our data suggests that Ras-mediated MAPK-ERK signaling may play a role in driving both cuSCC and hepatocellular adenocarcinoma. In support of this, the Onc2.3-driven cuSCCs also exhibited inactivation of *Nf1*, encoding a negative regulator of Ras signaling important in many cancer types. Mutations in *RASA1* and *NF1* co-occur in human non-small cell lung cancer (Hayashi et al.), which may reflect independent mechanisms by which RASA1 and NF1 inhibit RAS or the clonal diversity within these tumors that preferentially selects one of these mechanisms.

The majority of the putative driver genes identified in HCA driven by Onc2.3 were also reported in SB-driven hepatocellular carcinoma (HCC)(Bard-Chapeau et al., 2014; Dupuy et al., 2009; Keng et al., 2013; O’Donnell et al., 2012; Rogers et al., 2013), supporting the role for these events in HCC initiation. While we did not identify SB inaction of *Ncoa2* and *Sav1* in our tumors, these genes were previously identified in liver cancer mouse models driven by *SB* mutagenesis in the presence of Myc activation or *Pten* loss (Kodama et al., 2018; O’Donnell et al., 2012). It is likely that initiating oncogenic events influence the evolutionary routes of tumor development, dictating the selection and maintenance of cooperating drivers. The extent to which context-dependent SB insertion events impact tumor development remains to be determined. Our HCA/HCC data suggests that while tumors initiated by an oncogenic mutation may exhibit shorter tumor latencies than tumors lacking exogenous initiating oncogenic events and may have distinct subsets of inactivated genes, both populations converge on similar signaling pathways. Importantly, the frequent development of cuSCC and HCA using the Onc2.3 SB transposon demonstrates that tumor suppressor gene inactivation is sufficient for *de novo* transformation of both keratinocytes or hepatocytes *in vivo*.

When all the Onc2.3 tumors were analyzed together, we noted that the negative regulators of the Ras signaling pathway such as *Rasa1* and *Nf1* were commonly inactivated in liver and cuSCC, while genes encoding members of the E3 ubiquitin ligase such as *Trip12* and *Cul3* were frequently inactivated in all tumor types analyzed. Additionally, in cuSCC tumors without activation of the *Zmiz* proto-oncogenes, the frequency of *Trip12* and *Cul3* inactivation were markedly increased, relative to tumors with *Zmiz* oncogene activation. The E3 ubiquitin ligase-proteasome have diverse roles, including regulating the Ras, MAPK, and PI3K/Akt signaling pathways, gene expression and cell death (Senft et al., 2018). This range of biological processes may explain why *Trip12* and *Cul3* inactivation is frequently observed in the Onc2.3 tumors. Indeed, a previous *Sleeping Beauty* screen by Dorr *et al*. using the Onc2 transposon system also found inactivation in *Cul3* (Dorr et al., 2015). The authors functionally validated the tumor suppressive role of *CUL3* in A549 and H522 human lung cancer cell lines, and showed that shRNA knockdown of this gene increased proliferation rates, relative to control. The second E3 ligase associated gene *Trip12* observed in our screen has been shown to be essential in the regulation of the cell cycle (Larrieu et al.).

Our Onc2.3 model is not without limitations. First, in cuSCC tumors, it is not clear why we did not observe inactivation of tumor suppressors frequently described in human squamous epithelium such as *TP53* and *NOTCH1*. Inactivation of *NOTCH1* has been postulated to drive the evolutionary dead-end of nascent pre-cancerous population of cells (Higa and DeGregori). This is based on the observation that the frequency of *NOTCH1* mutations decreases in esophageal and skin cancers, relative to physiologically normal epithelial tissues (Higa and DeGregori, 2019; Martincorena et al., 2018; Yokoyama et al., 2019). Consistent with this, in our previously reported Onc3 screen, we noted that the frequency of *Notch1* inactivation was decreased in cuSCC versus keratoacanthoma tissues, suggesting that *Notch1* loss is not essential for cuSCC development (Aiderus et al.). While we did not observe inactivation of *Trp53*, our previous Onc3 screen suggests that there were no obvious differences in the insertion profiles of tumors with either mutant or wild type *Trp53* background, albeit the rate of tumorigenesis was significantly reduced in *Trp53* mutant mice. Additionally, as noted in the SB Cancer Driver Database, the low frequency of transposon-driven *Trp53* inactivation is not unique to our study, and has been observed in other SB screens reported to date (Newberg et al.). Second, we noted that genes involved in chromatin remodeling were inactivated in all of the tumor types sequenced, albeit at different frequencies. It is possible that loss of these tumor suppressors may activate oncogene expression, thereby promoting tumorigenesis. However, given the significantly extended latency of tumor development in the Onc2.3 mice, it is unlikely that such genes confer strong advantage in clonal expansion, and the chromatin remodeler loss likely affects tumor suppressive processes. Similarly, without genomic sequencing analysis, we cannot exclude the possibility that spontaneous driver mutations that confer cooperating oncogenic and/or tumor suppressive alterations, are also contributing to the Onc2.3-driven tumors analyzed in this study. However, extensive genomic analyses of whole genome, exome or transcriptome from SB-driven melanoma(Mann et al., 2015; Perna et al., 2015), myeloid leukemia (Mann et al., 2016b) and cuSCC (Aiderus et al., 2019) have revealed very few non-transposon derived alterations, including any known or suspected oncogenic events. Last, in this study, we focused only on determining the transposon insertion profiles across various tumor types. The inactivation of genes involved in chromatin remodeling and protein ubiquitination suggests the necessity to profile the transposon mutagenesis-driven tumors on other platforms (*e.g*., transcriptome, proteome) to gain a more comprehensive understanding of the molecular events driving tumorigenesis.

Collectively, our data strongly implicate routes to tumorigenesis via cooperative inactivation of tumor suppressor driver genes alone, and precludes the necessity of oncogenic events. The dramatic reduced penetrance and increased latency of skin, lung, and liver tumors when subjected to systemic gene-trap only transposon mutagenesis, highlights three important points: i) gene inactivation *per se* is sufficient to drive *de novo* transformation *in vivo* of keratinocytes, lung alveolar epithelium, and hepatocytes and tumor initiation *in vivo*; ii) transformation by gene inactivation alone is likely to be the exception rather than rule as penetrance of proto-oncogenic driven cuSCC, LUAC, and HCA is substantially higher, and iii) cuSCC development has no absolute requirement for oncogenic drivers (*e.g*., activating ZMIZ or RAS oncoproteins), and can occur via cumulative inactivation of tumor suppressors, which in either instance, converge on similar altered cancer hallmark signaling pathways. Thus, we conclude that the SB-Onc2.3 model delivers a novel *in vivo* platform that allows systemic gene inactivation in a non-sensitized background, provides a heterogeneous background that promotes evolutionary routes to tumorigenesis.

## Author Contributions

M.B.M., K.M.M. N.G.C., and N.A.J. conceived the project and provided laboratory resources and personnel for animal husbandry, specimen archiving, sequencing and compute management. M.B.M. and K.M.M. designed the studies, directed the research, interpreted the data, performed experimental work, coordinated sequencing, and analyzed the data. A.A., A.M.C.-S., and A.L.M. performed and/or analyzed experimental datasets, including optimizing capture hybridizations for the SBCapSeq experiments. J.Y.N. and M.B.M. performed statistical and bioinformatics analyses. J.M.W. performed and directed veterinary pathology analysis, including tumor grading and diagnosis. D.S. generated the high-copy T2/Onc2.3 transgenic founder mouse lines. A.A., K.M.M. and M.B.M. wrote the manuscript. All co-authors approved manuscript before submission.

---

**Supplementary Figure 1:**
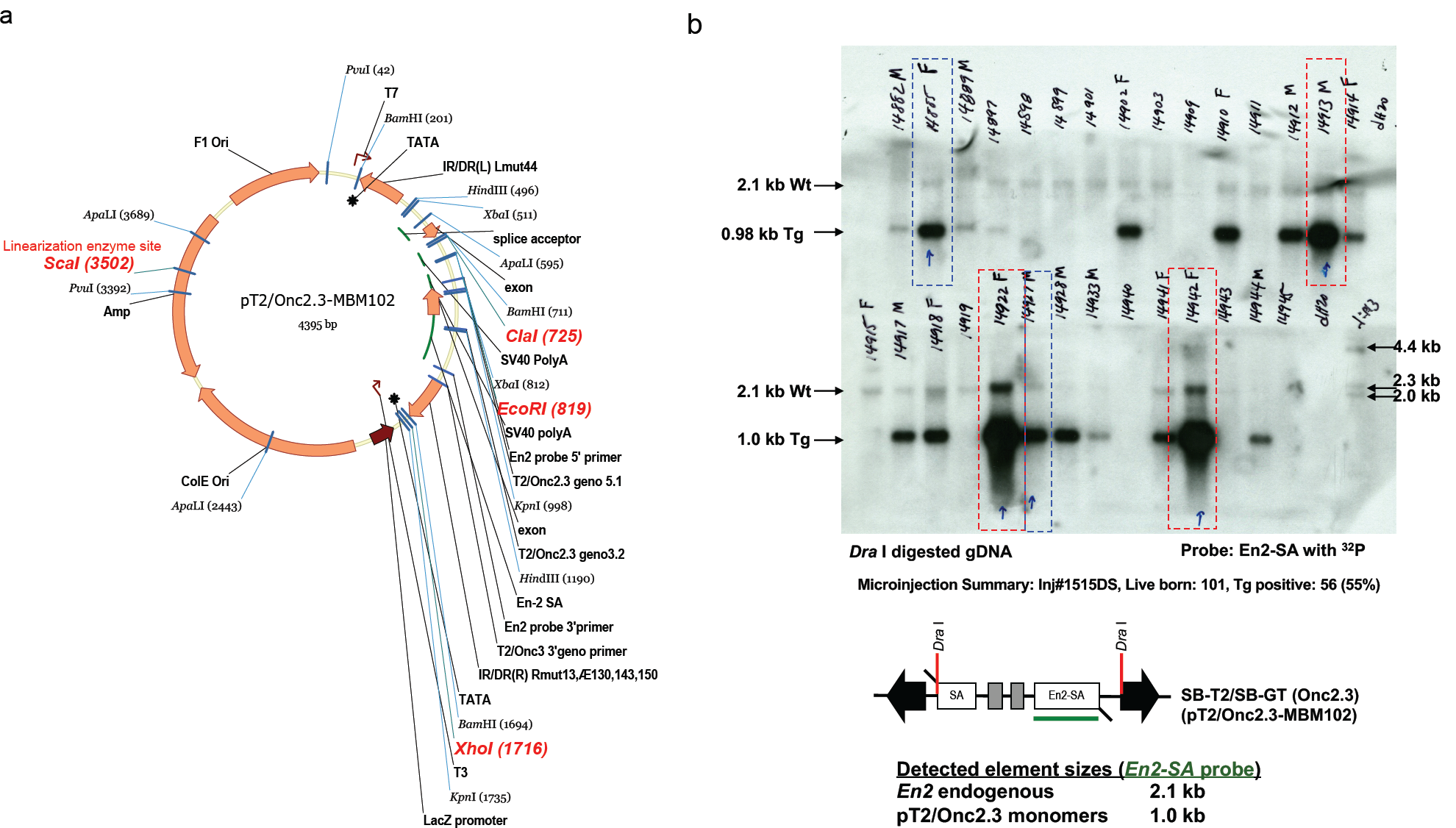
Transgenic SB-T2/Onc2.3 founder mice. (**a**) Plasmid map of pT2/Onc2.3-MBM102, denoting essential features and base pair position in parenthesis, used to create the T2/Onc2.3 high-copy transgenic lines. (**b**) Southern blot of T2/Onc2.3 founder mice using a probe corresponding to the *En2*-SA (*Engrailed 2* splice acceptor) element within the SB-T2/SB-GT allele construct that also cross hybridizes to the endogenous *En2* mouse gene on chromosome 5. Genomic DNA digested to completion with *Dra*I restriction enzyme and detected with a radioisotope-^32^P labeled *En2*-SA probe identifies the diploid endogenous *En2* locus as a discrete 2.1 kb band and any all copies of the T2/SB-GT monomer that have been liberated from the their randomly inserted transgenic loci as a discrete 1.0 kb band. Mice with multi-copy concatemer alleles are identified as containing darker staining bands relative to the two copies of the *En2* endogenous locus band. Blue arrows and dotted boxes denote animals selected for germ line breeding — five separate transgenic founders, three females (TG.14885, TG.14922, and TG.14942) and two males (TG.14913 and TG.14927), were selected for mating with wildtype C57BL/6J mice to confirm the germ line transmission of the SB-T/2SB-GT concatemer alleles to progeny. Mice denoted with blue dotted boxes produced transgenic carrier pups with a transgene copy numbers that differed between individuals, suggesting the presence of two or more independent transgenic concatemer integration sites. Mice denoted with red dotted boxes produced transgenic carrier pups with a transgene copy number that did not differ between individuals, suggesting that they resulted from a single transgenic concatemer integration site. Lines established from founders TG.14913, TG.14922, and TG.14942 were expanded and bred for further experiments. *Bam*HI restriction digested *lambda* (ƛ)-phage DNA is provided as a reference, a small amount was added to the radioisotope-^32^P labeling reaction with the *En2*-SA probe to show detection on the developed Southern blot.

**Supplementary Figure 2:**
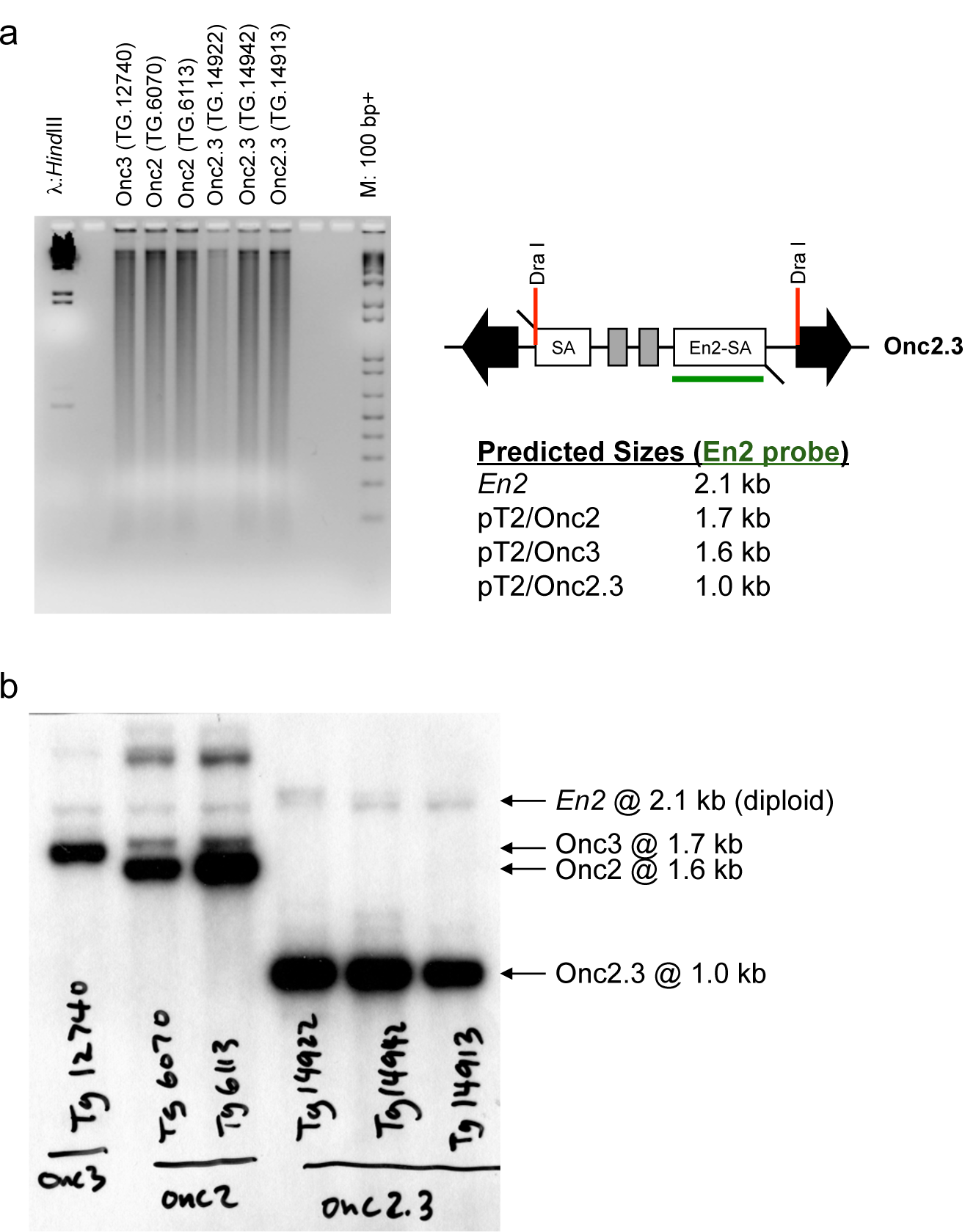
Estimating SB transposon concatemer copy number from different SB strains. (**a**) Inverse image of EtBr stained agarose gel (1.2% TBE, run at 30 volts for 30 hours) containing genomic DNA digested with *Dra* I for 16 hours at 37 °C for Southern blot analysis. Molecular weight markers: left, lambda cut with *Hin* DIII; right, 100 bp ladder. (**b**) Southern blot demonstrating relative concatemer copy number differences between high-copy and low-copy SB transposon alleles in homozygous mice with unmobilized transposons. The *En2* site provides a reference for a single copy, diploid gene.

**Supplementary Figure 3:**
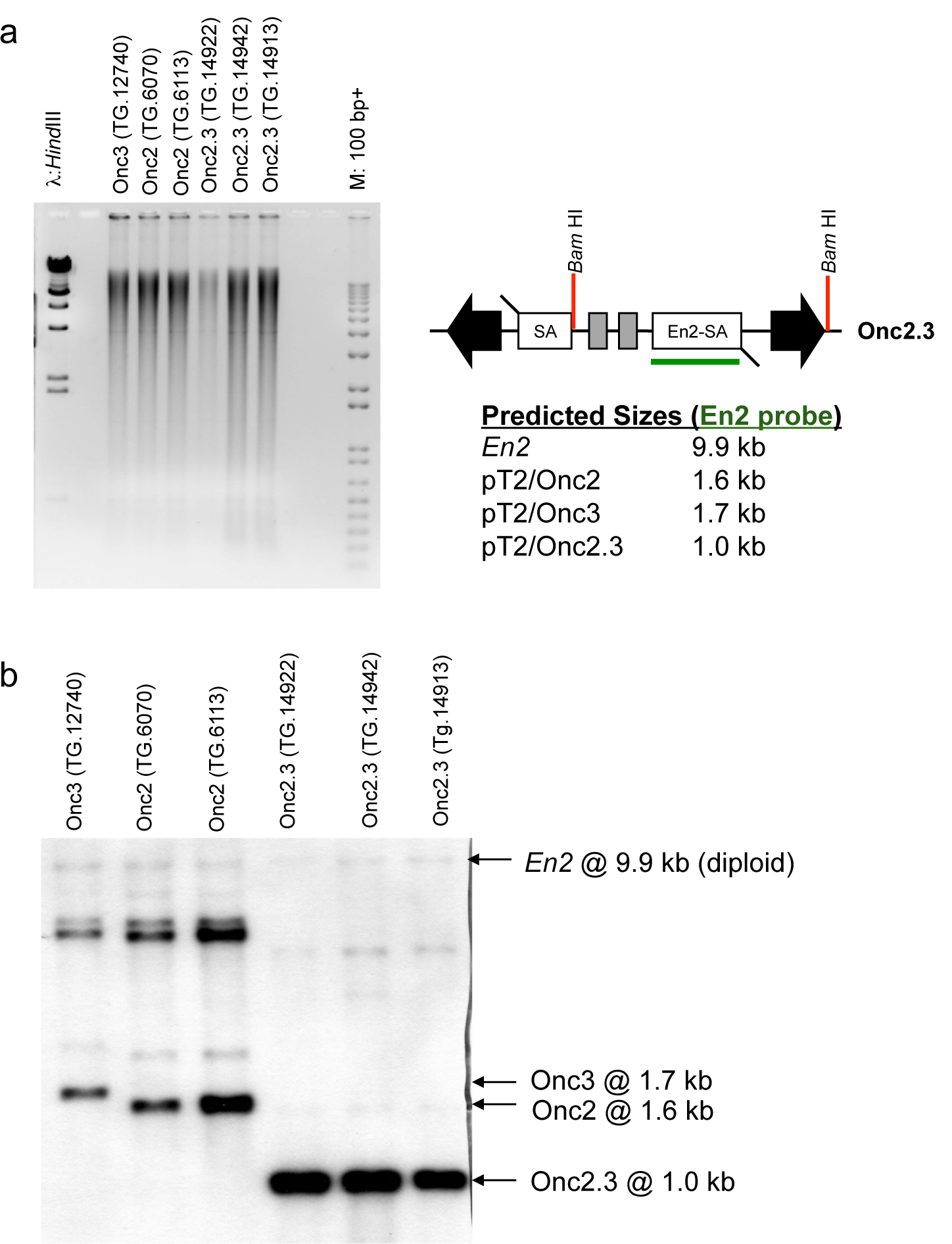
Estimating SB transposon concatemer copy number from different SB strains. (**a**) Inverse image of EtBr stained agarose gel (1.2% TBE, run at 30 volts for 30 hours) containing genomic DNA digested with *Bam*HI for 16 hours at 37 °C for Southern blot analysis. Molecular weight markers: left, lambda cut with *Bam*HI; right, 100 bp ladder. (**b**) Southern blot demonstrating relative concatemer copy number differences between high-copy and low-copy SB transposon alleles in homozygous mice with unmobilized transposons. The *En2* site provides a reference for a single copy gene.

**Supplementary Figure 4:**
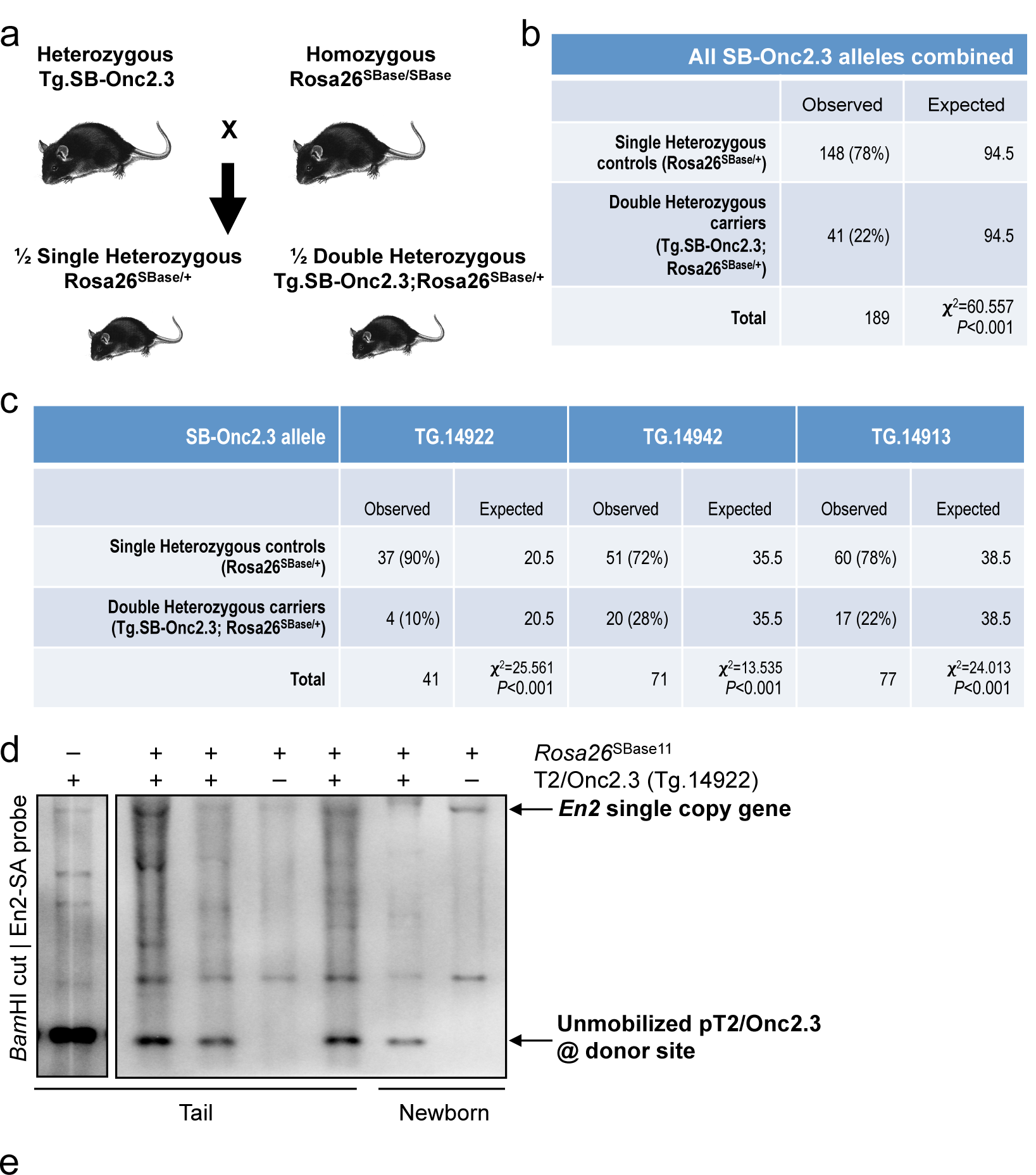
Embryonic lethality in double heterozygous mice. (**a**) Mouse cross to generate singe allele control and double allele experimental mice. (**b**) Less than expected number of progeny resulted from matings between double heterozygous carrier mice resulting from crosses between SB T2/Onc2.3 and constitutive SBase expression. (**c**) Onc2.3 (Tg/+); SB’B’ (neo/+) double hets are not born in the expected Mendelian frequencies, Goodness of Fit test values provided. (**d**) SB transposition and genome mobilization of T2/Onc2.3 allele *in vivo* by constitutive SBase expression in compound double heterozygous carriers. Southern blot demonstrating change in T2/Onc2.3 (Tg.14922) allele donor site concatemer copy number of in single carrier mice with unmobilized transposons (left lane) and significant depletion in double carrier mice from tail biopsies of wean pups or from whole body newborn kidney specimen. (**e**) Modified SB excision PCR(Mann et al., 2015) demonstrating transposition of the T2/Onc2.3 (Tg.14922) allele *in vivo* from routine tail biopsies taken from weaned mice for genotyping. Presence of the SB excision PCR product infers SB transposon and SB transposase alleles are all present within the germ line of the weaned mouse.

**Supplementary Figure 5:**
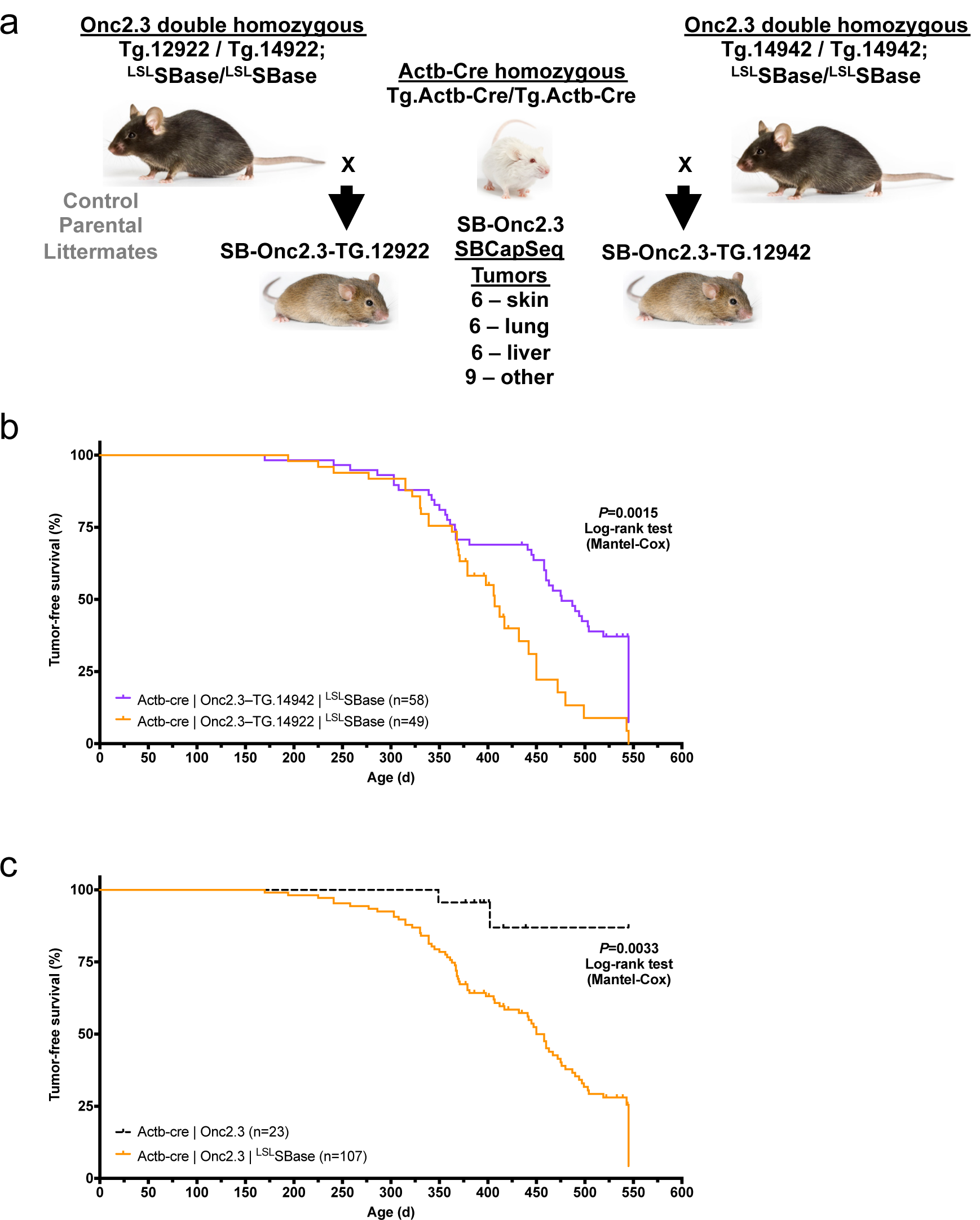
Overview of SB-induced tumor studies. (**a**) SB-Onc2.3 breeding strategy and genotype cohorts aged to generate the SB-Onc2.3 experimental and control cohorts of mice used in this study. (**b**) Kaplan-Meier survival plots comparing SB-Onc2.3 cohorts (Mantel-Cox log-rank test, *P* = 0.0015). (**c**) Kaplan-Meier survival plots comparing the combined SB-Onc2.3 experimental and control cohorts (Mantel-Cox log-rank test, *P* = 0.0033).

**Supplementary Figure 6:**
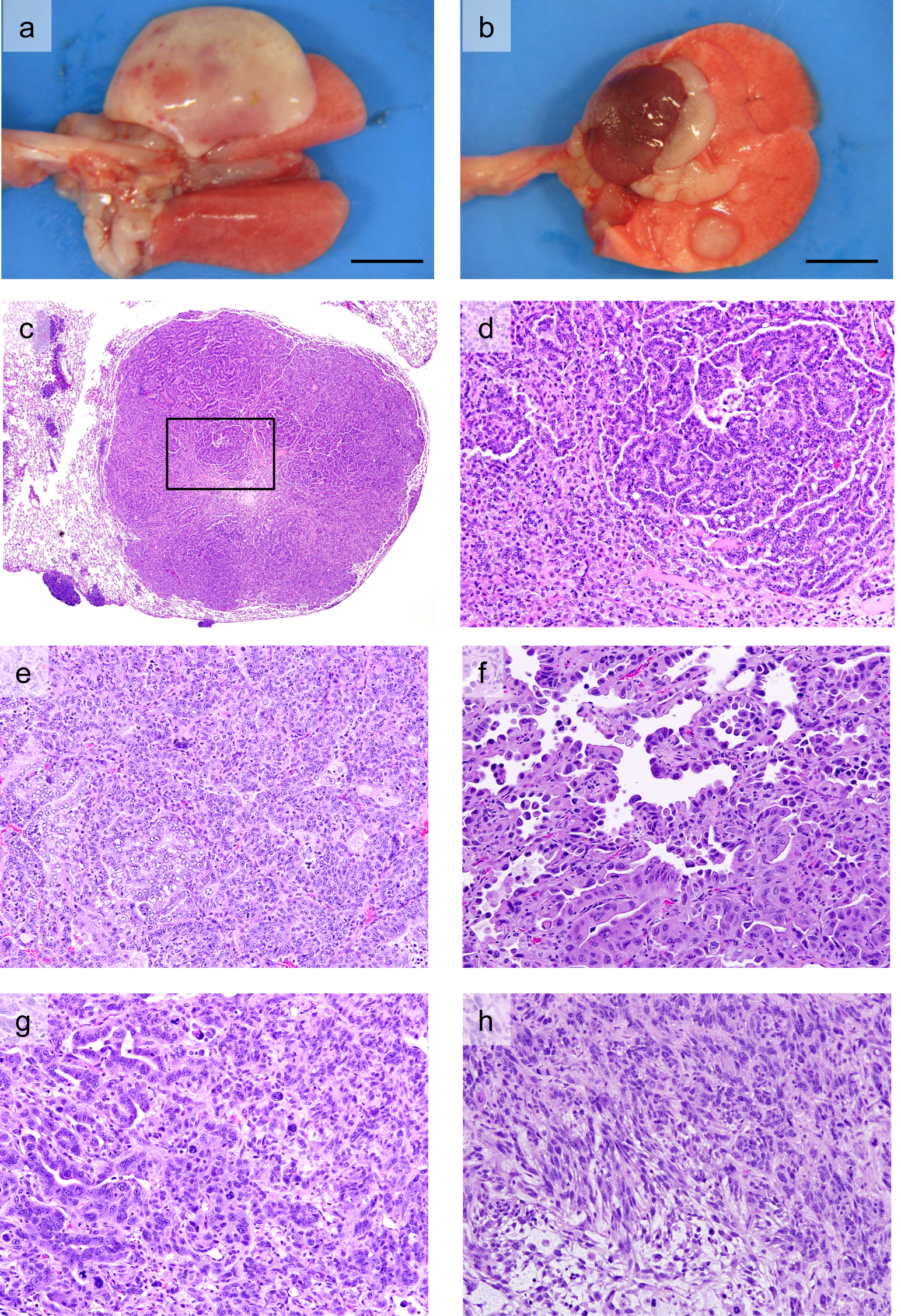
Necropsy and histological analysis of SB-Onc2.3 tumors. (**a-b**) Necropsy image of lung tumors: (**a**) white mass (adenocarcinoma) from right apical lobe and (**b**) small white nodule at bottom (adenoma) and at top of lung - large white mass with red area (adenocarcinoma) from the right ventral lobe. Histology and tumor classification from sections of lung masses stained with hematoxylin and eosin (H&E): (**c-d**) Lung adenoma (40×), with inset showing carcinoma differentiation in (**d**) (200×) and Lung (**e-h**) adenocarcinoma areas 200X, (**e**) carcinoma (200×), (**f**) adenocarcinoma (200×), and (**g**) sarcoma (200×). Source image files and additional histology images may be downloaded from figshare http://dx.doi.org/10.6084/m9.figshare.12816305.

**Supplementary Figure 7:**
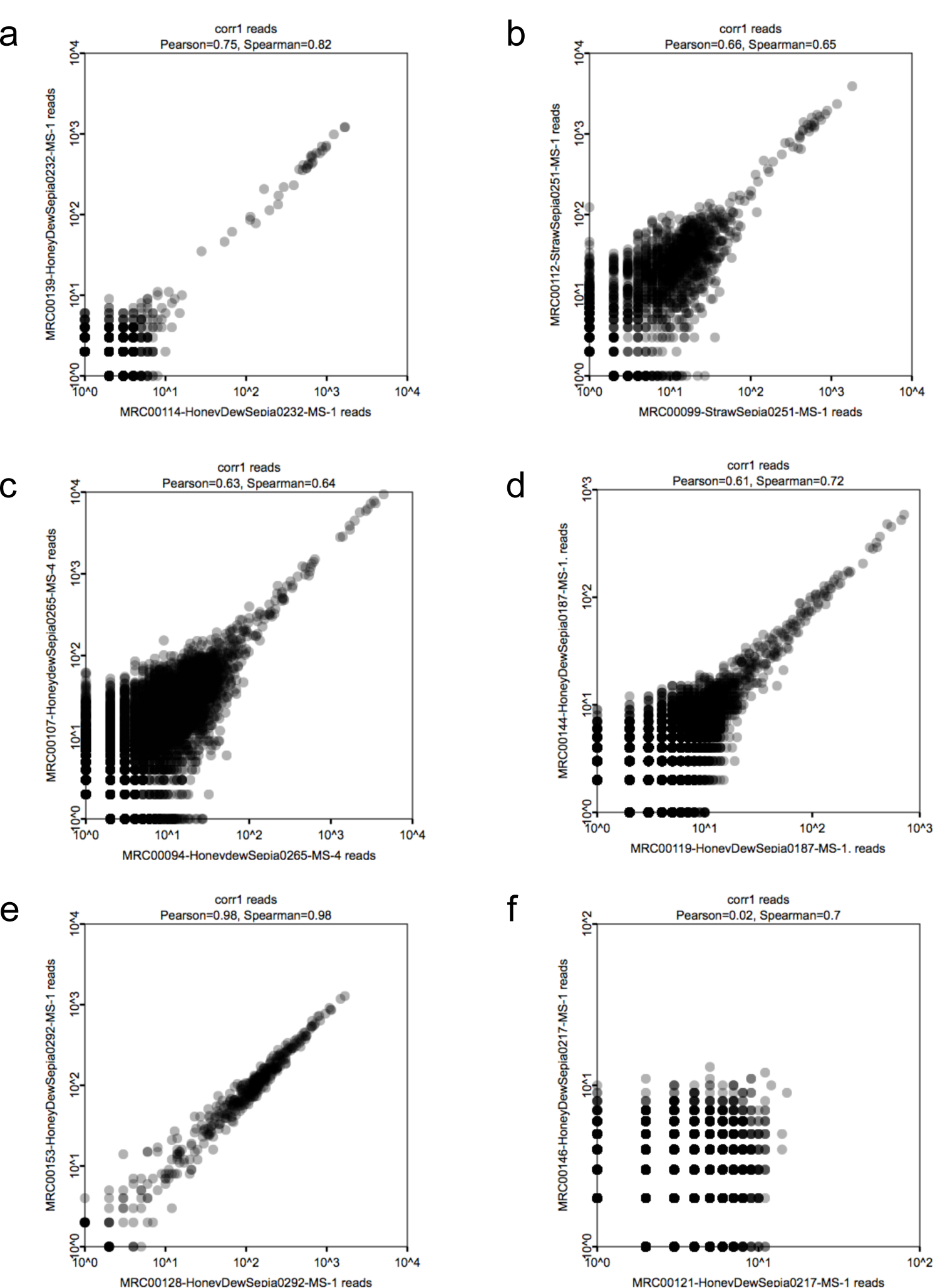
Evaluating the reproducibility of SBCapSeq from bulk SB-Onc2.3 tumors. Representative plots of the individual SB insertion sites based on read depth from (**a**) cuSCC, (**b**) HCA, (**c**) LUAA, (**d**) Ovarian adenocarcinoma, (**e**) Schwannoma, and (**f**) astrocytoma genomes from genomic DNAs created from bulk tumor specimens from technical replicate libraries (independent library sequencing runs applied to the same biological specimen isolate) comparing library run 1 (*x*-axis) to library run 2 (*y*-axis). Biological reproducibility of the cuSCC, HCA, LUAA, ovarian adenocarcinoma, and Schwannoma specimen libraries is high, supported by both Pearson’s and Spearman’s correlation metrics, and identifies the all of SB insertion sites with read depths of 20 or higher (**a-e**). In contrast, technical reproducibility of the astrocytoma specimen libraries is exceptionally low and fails to identify any read depth insertion sites higher than 22 reads (**e**), indicating that the insertions are likely background passenger insertion events that are not clonally selected during astrocytoma progression.

**Supplementary Figure 8:**
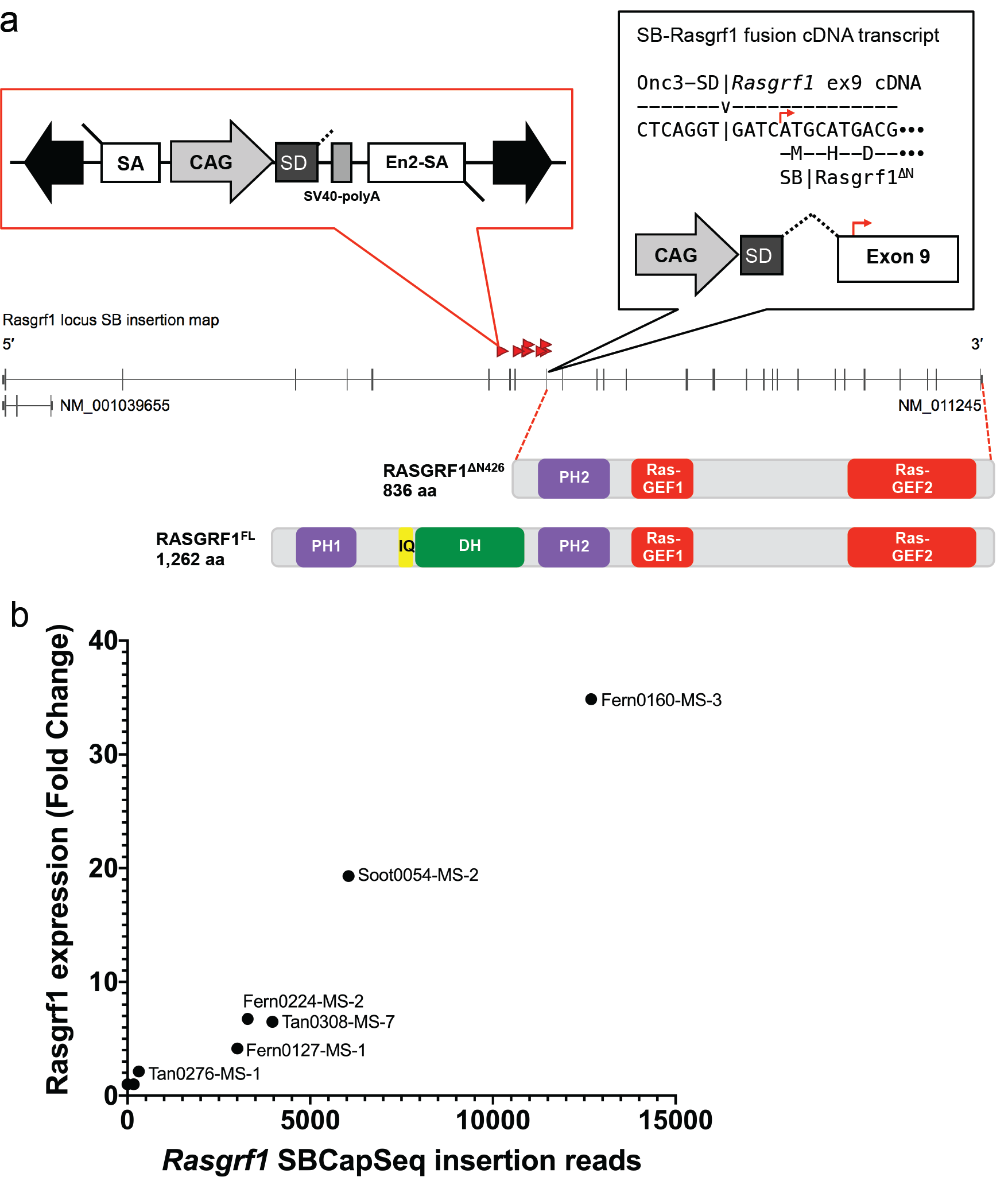
Activating SB insertions into the *Rasgrf1* locus drive overexpression an oncogenic delta-N isoform. (**a**) Recurrent activating SB insertions in Onc3-induced lung cancer genomes drives expression of an N truncated RASGRF1^ΔN426^ isoform that lacks the first 426 amino acids in the full length RASGRF1 protein. The protein domains for RASGRF1 from UniProt (http://www.uniprot.org/uniprot/P27671): PH1 (aa 22 – 130 in the full length), IQ (aa 208 – 233 in the full length), and DH (aa 244 – 430 in the full length) absent in the predicted RASGRF1^ΔN426^ oncoprotein; PH2 (aa 460 – 588 in the full length), Ras-GEF1 (aa 635 – 749 in the full length) and Ras-GEF2 (aa 1027 – 1259 in the full length) present in the predicted RASGRF1^ΔN426^ oncoprotein. (**b**) Expression of the delta-N *RASGRF1* is positively correlated with SBCapSeq read depth (Pearson r=0.9761; Spearman r=0.9848; *P*<0.0001; **Supplementary Table 24**).

**Supplementary Figure 9:**
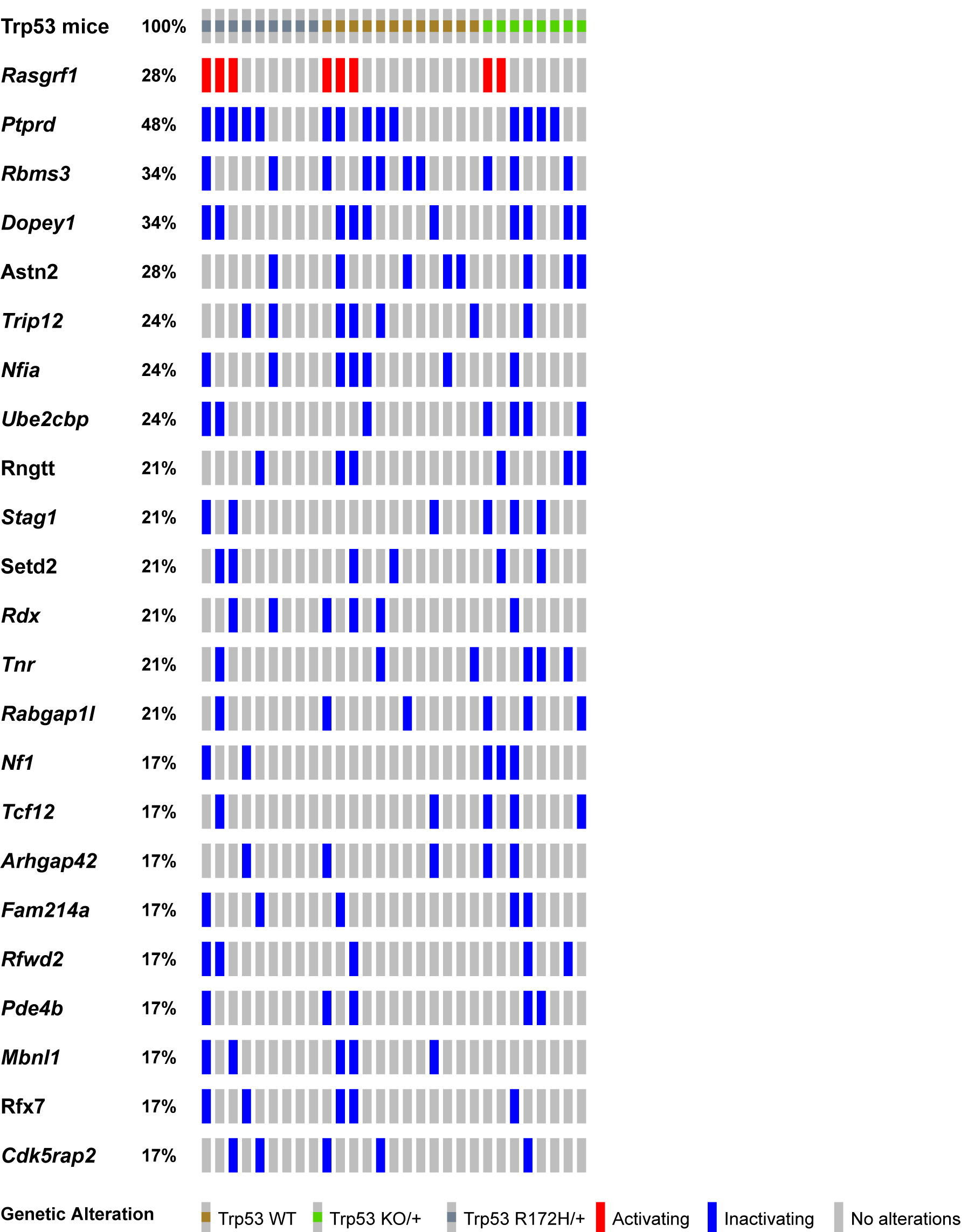
Landscape of candidate trunk drivers mutated in SB-induced LUAA with Splink_454T sequencing. SB Driver Analysis applied to Splink_454 SB insertion data (**Supplementary Table 8**) with read depth cutoff of 6 on 29 late-stage HCA genomes. FWER-corrected significant driver genes (rows) were sorted by occurrence within the cohort genomes (columns).

**Supplementary Figure 10:**
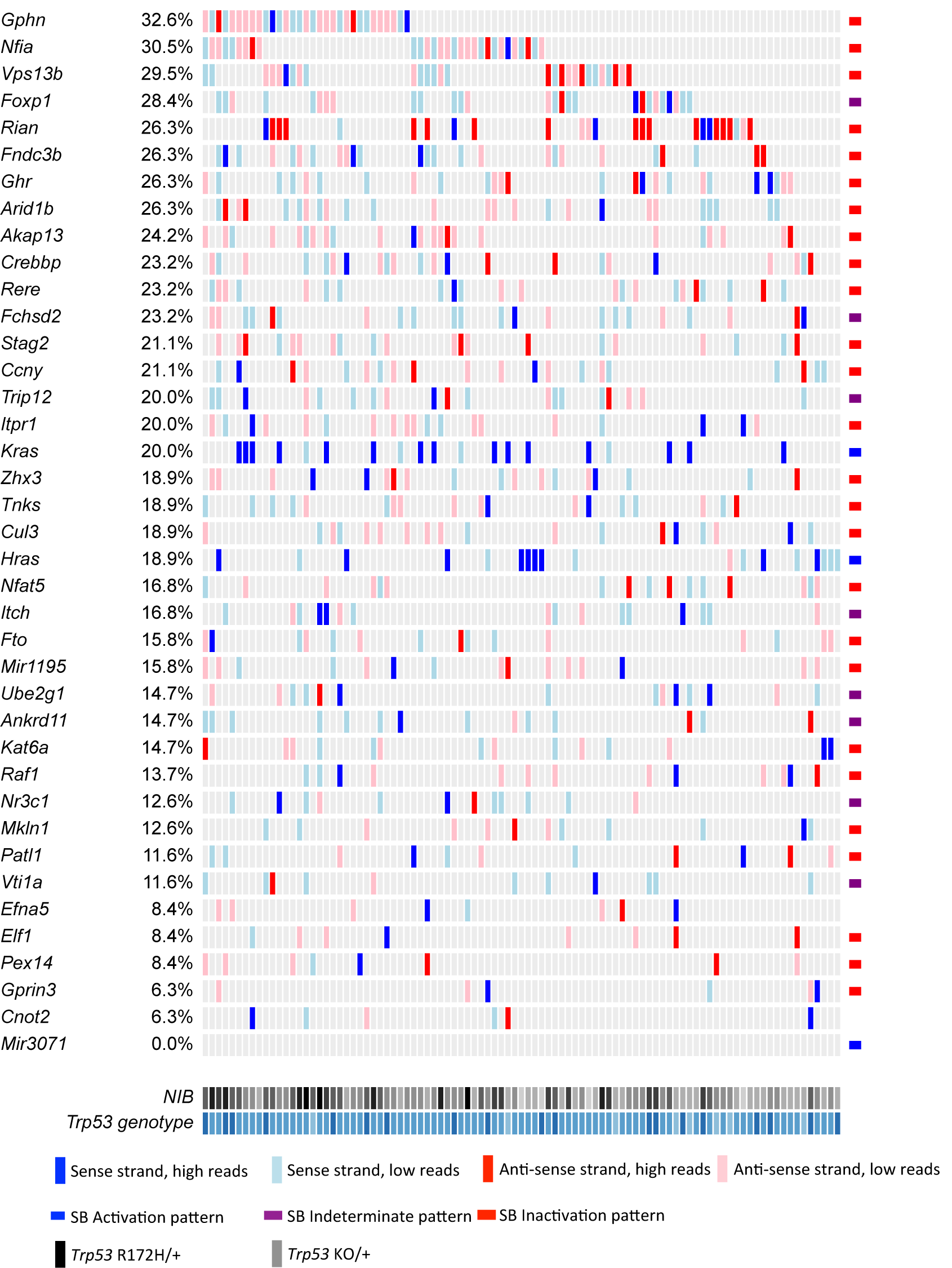
Landscape of candidate trunk drivers mutated in SB-induced HCA with Splink_454T sequencing. SB Driver Analysis applied to Splink_454 SB insertion data (**Supplementary Tables 10-12**) with read depth cutoff of 6 on 95 late-stage HCA genomes. FWER-corrected significant driver genes (rows) were sorted by occurrence within the cohort genomes (columns). Waterfall plots were generated on all insertions, and insertion patterns determined by progression driver analysis are shown to the side. NIB, normalized insertion burden from lowest (light gray) to highest (black). *Trp53* genotype shown from *Trp53* ^+/+^ (light blue), *Trp53* ^R172H/+^ (blue) and *Trp53* ^KO/+^ (dark blue).

## Methods

### Generation of pT2/Onc2.3 vector

SB transposon vector pT2/Onc2(Dupuy et al., 2005) (a gift from A. Dupuy) was sequentially digested with restriction enzymes *Xma*I (NEB), at position 717 bp, and *Eco*RI (NEB) at position 729 bp, at 37 °C for 30 m each, per manufacturer’s suggested protocol, to linearize the plasmid. Linear plasmid was visually confirmed as an ∼4.9 kb on a 1% agarose TAE gel stained with EtBr viewed under UV light. A small piece of agarose gel containing the linear pT2/Onc2 was excised, purified using GFX columns (Amersham Biosciences #414581), and saved for ligation. A PCR cloning strategy was used to insert an extra bi-directional SV40-polyA termination sequence into the pT2/Onc2 vector. Using 100 uM each of two complimentary overlapping (**bold**) oligos, <u>XmaI</u>+ClaI.SV40-pA 5’-ATG CAT GCA T<u>CC</u> <u>CGG G</u>CA TCG ATG CAG TGA AAA AAA TGC TTT ATT TGT GAA ATT T**GT GAT GCT ATT GCT TTA TTT GTA ACC**-3’ and SV40-pA.<u>EcoRI</u>+XbaI 5’-ATG CAT GCA TGA ATT CGT CTA GAA ACT TGT TTA TTG CAG CTT ATA AT**G GTT ACA AAT AAA GCA ATA GCA TCA C**-3’ and PCR SuperMix Taq (Invitrogen #10572014), we amplified a 128 bp fragment (5’-ATG CAT GCA T<u>CC CGG G</u>CA TCG ATG CAG TGA AAA AAA TGC TTT ATT TGT GAA ATT TGT GAT GCT ATT GCT TTA TTT GTA ACC ATT ATA AGC TGC AAT AAA CAA GTT TCT AGA CGA ATT CAT GCA TGC AT-3’) using at thermal cycler program: template denaturation at 94°C for 90 s followed by 30 cycles of 94°C for 30 s, 50°C for 45 s, and 72°C for 45 s followed by 72°C for 300 s and cooled to 4°C. PCR product was visually confirmed by running the PCR product on a 2% agarose TBE gel stained with EtBr viewed under UV light. A small piece of gel containing the linear PCR product was excised, purified using GFX columns (Amersham Biosciences #414581), and used to ligate into pGEM^®^-T Easy Vector System (Promega #A1360) at room temperature for 60 m using the manufacturer’s suggested protocol. Ligation products were electroporated into MAX efficiency DH10B competent cells (Invitrogen #18297010), plated on LB agar plates with ampicillin, and incubated 37 °C for 18 hrs. Miniprep DNA from selected transformants was sequentially digested with *Xma*I (NEB) and *Eco*RI (NEB), at 37 °C for 30 m each, per manufacturer’s suggested protocol. The ligation product was run on a 2% agarose TBE gel stained with EtBr viewed under UV light, and a small piece of gel containing the 108 bp fragment was excised, purified using GFX columns (Amersham Biosciences #414581) prior to ligation. Finally, a ligation reaction using a 1:5 ratio of pT2/Onc2:XmaI-EcoRI linear vector and SV40-polyA: :*Xma*I-*Eco*RI insert, respectively, was performed using T4 DNA Ligase at room temperature for 30 m, followed by heat inactivation at 65 °C for 20 m. Ligation products were electroporated into MAX efficiency DH10B competent cells (Invitrogen #18297010), plated on LB agar plates with ampicillin, and incubated at 37 °C for 18 hrs. Miniprep DNA from selected transformants was digested with *Hin*dIII (NEB), at 37 °C for 60 m each, per manufacturer’s suggested protocol. Recombinant pT2/Onc2.2 vector was identified by the presence of three fragments at 3.658 kb, 0.8 kb, and 0.6 kb fragments. pT2/Onc2.2 vector DNA was cut with restriction enzyme AscI (NEB #R0558S), at 37 °C for 60 m, to linearize the plasmid. Linear plasmid was visually confirmed on a 1% agarose TAE gel stained with EtBr viewed under UV light. A small piece of agarose gel containing the linear pT2/Onc2.2 vector was excised and purified using GFX columns (Amersham Biosciences #414581). PCR amplification was performed using 100 ng of the purified linearized pT2-Onc2.2 vector template and a pair of oligos T2/Onc2.f1 5’-ATG C<u>GA ATT C</u>AA CGC GCG TTA AGA TAC ATT GA-3’ engineered to contain an <u>EcoRI</u> site (underlined) and T2/Onc2.r1 5’-ATC TAT GGC TCG TAC TCT ATA GGC-3’ designed to amplify the pT2/Onc2 vector excluding the MSCV 5’ LTR minimal promoter and Lun-SD sequences using platinum Hi-Fi DNA *Taq* polymerase (Invitrogen #11304-028) in 1X Hi-Fi PCR buffer (Invitrogen #52045) supplemented with 2 uL of 50 mM MgSO4 and dNTPs (Roche#11-581-295-001) and thermal cycler program: template denaturation at 95°C for 90 s followed by 30 cycles of 95°C for 45 s, 55°C for 60 s, and 72°C for 270 s followed by 72°C for 600 s and cooled to 37°C. Then, 1 unit of restriction enzyme *Dpn*I (NEB #R0176S) was added (to digest the methylated plasmid from vector template grown in bacteria and enrich PCR amplified template during ligation) to each PCR tube and incubated at 37°C for 30 m, followed by heat inactivation at 80°C for 20 m, and then cooled to 4 °C. The PCR product was column purified, resuspended in ddH2O, and digested with *Eco*RI (NEB) at 37 °C for 60 m (one EcoRI site was added by the oligo and the other occurs within the pT2/Onc2.2 vector at position 819). Gel and column purified pT2/Onc2.3 linear vector (4.395 kb) was self-ligated using T4 DNA ligase at room temperature for 30 m, followed by heat inactivation at 65 °C for 20 m. Ligation products were electroporated into MAX efficiency DH10B competent cells, plated on LB agar plates with ampicillin, and incubated at 37 °C for 18 hrs. Miniprep DNA from selected transformants was digested with *Hin*dIII (NEB), at 37°C for 60 m each, per manufacturer’s suggested protocol. Recombinant pT2/Onc2.3 vector (**Supplementary Figure 1a**) was identified by the presence of two fragments at 3.701 kb and 0.694 kb fragments. Cloning maps and sequence files may be downloaded from figshare http://dx.doi.org/10.6084/m9.figshare.12816452. Several pT2/Onc2.3 vector clones were sequenced using BigDye^®^ direct cycle sequencing kit, using the manufacturer’s suggested protocol, and the following primers: T7, 5’-GTAATACGACTCACTATAGGG-3’; T2/Onc.sp5.1, 5’-GACTGTGCCTTTAAACAGCTTGG-3’; T2/Onc.sp5.2, 5’-TCCTGTGCCAGACTCTGGCGC-3’; and T2/Onc.sp5.3, 5’-GGGTGGTGATATAAACTTGAGGCTG-3’. A single pT2/Onc2.3 clone with full sequence identity to the vector map was re-electroporated into MAX efficiency DH10B competent cells (Invitrogen #18297010), plated on LB agar plates with ampicillin, and incubated at 37°C for 18 hrs.

### Generation of T2/Onc2.3 transgenic mice

pT2/Onc2.3 DNA was linearized with restriction enzyme with *Sca*I (NEB #R3122) at position 3,502 bp (**Supplementary Figure 1a**) at 37°C for 90 m, followed by heat inactivation at 80 °C for 20 m. pT2/Onc2.3:*Sca*I linear plasmid, at 2 ng/uL, 4 ng/uL and 10 ng/uL, was prepared for microinjection into (B6C3)F2 hybrid embryos using standard techniques (Dupuy et al., 2005). Tail biopsy genomic DNA from founder animals was digested with *Dra*I (NEB) and *Bam*HI (NEB), run on a 0.8% TAE agarose gel at 30 v for 16 h, transferred to membrane, and screened by Southern blotting using a ^32^P-labeled En2-SA (splice acceptor) probe, a 500 bp En2-SA PCR product with pT2/Onc2.3 template and primers T2.3-En2.5Probe, 5’-GCTGCAATAAACAAGTTGGCCG-3’ and T2.3-En2.3Probe, 5’-CTTGGGTCAAACATTTCGAGTAGCC-3’, and standard methods. Individual transgenic lines (n=5) were established by backcrossing founders to C57BL/6J (JAX#000664). Germ line transmission was confirmed by Southern blotting using standard techniques. All five lines transmitted to offspring, but only three lines, T2/Onc2.3(TG.14913) (TgTn(sb-T2/Onc2.3)14913Mbm), T2/Onc2.3(TG.14922) (TgTn(sb-T2/Onc2.3)14922Mbm), and T2/Onc2.3(TG.14942) (TgTn(sb-T2/Onc2.3)14942Mbm), segregated a transposon concatemer as a single, discreet locus by Southern blot genotyping. Subsequent offspring were genotyped by multi-plex PCR using primers specific to the SB transposon T2.3-sense, 5’-CTG TCA GGT ACC TGT TGG TCT GAA AC-3’ and T2.3-antisense, 5’-CCT CAA GCT TGG GTG CGT G-3’, which yields a 370 bp PCR product, and control primers Actb-sense, 5’-ACA AGG TCA AAA CTC AGC AAC AAG T-3’ and Actb-antisense, 5’-GCT GAG AGG GAA ATT GTT CAT TAC A-3’, which yields a 700 bp PCR product. Multiplex PCR genotyping protocol for SB-Onc2.3 mice may be downloaded from figshare http://dx.doi.org/10.6084/m9.figshare.12811301.

The following alleles were used to construct the SB-driven mouse model of multiple solid tumor histologies: *Actb-Cre* (FVB/N-Tg(ACTB-cre)2Mrt/J, ref. (Lewandoski et al., 1997));*Trp53*^*flox/+*^ (FVB.129P2-*Trp53*^*tm1Brn*^/Nci, ref. (Jonkers et al., 2001)); *Trp53*^*LSL-R172H/+*^ (129S4-*Trp53*^*tm2Tyj*^/Nci; ref. (Olive et al., 2004)); T2/Onc2(TG.12740) (TgTn(sb-T2/Onc3)12740Njen; ref. (Dupuy et al., 2009)); and *Rosa26*-LSL SBase or SBase^LSL^; (Gt(ROSA)26Sor^tm2(sb11)Njen^; ref. (Starr et al., 2009)). The resulting cohorts of mice were on mixed genetic backgrounds consisting of C57BL/6J, 129, C3H and FVB. Genotyping by PCR assays with primers specific to the alleles was performed. No sample size estimate was used. The production and characterization of all SB and SBase mouse strains was supported by the Department of Health and Human Services, National Institutes of Health and the National Cancer Institute. Mice were bred and maintained in accordance with approved procedures directed by the respective Institutional Animal Care and Use Committees in the National Cancer Institute Frederick National Lab, A*STAR Biological Resource Centre, and H. Lee Moffitt Cancer Center and Research Institute/University of South Florida. Gross necropsies were performed and all masses were documented and prepared for subsequent analysis. Both sexes were used for experiments, including reporting of cohort data survival analysis. No randomization or blinding was performed, all mice were assigned to groups based on genotype.

### Histological analysis

Histological analysis of necropsy tissues was performed on 5-μm sections of formalin-fixed, paraffin-embedded (FFPE) samples stained with hematoxylin and eosin (H&E). Nuclear staining of SB transposase (SBase) was confirmed by immunohistochemistry on FFPE tissues after antigen retrieval (pH 9) and endogenous peroxidase inhibition followed by overnight incubation with mouse antibody to SBase (anti-SBase; R&D Systems; pH 9; 1:200 dilution), followed by incubation with primary antibody, chromogen detection (with HRP polymer, anti-rabbit or anti-mouse, with Envision System from Dako) and hematoxylin counterstaining were performed per manufacturer’s instructions. Genomic DNA (gDNA) was isolated from flash frozen or FFPE necropsy specimens using Qiagen Gentra^®^ Puregene^®^ DNA isolation kit protocol for tissue using the manufacturer’s protocol (see **Supplementary Note 2**).

### Mapping SB insertion sites

Full details for the SBCapSeq protocol, an enhanced SBCapSeq method optimized for sequencing from solid tumors (Mann et al., 2019), will be published elsewhere (Mann, Mann *et al*., in preparation; for general protocol and concept, see also refs. (Mann et al., 2016a; Mann et al., 2016b)). Briefly, for selective SB insertion site sequencing by liquid hybridization capture, gDNA (0.5 µg per sample) of bulk tumor genomes was used for library construction using the AB Library Builder(tm) System, including random fragmentation and ligation of barcoded Ion Xpress(tm) sequencing adapters.

Adapter-ligated templates were purified by Agencourt^®^ AMPure beads^®^ and fragments with insert size of ∼200 bp were excised, purified by Agencourt^®^ AMPure^®^ beads, amplified by 8 cycles of adapter-ligation-mediated polymerase chain reaction (aLM-PCR), and purified by Agencourt^®^ AMPure^®^ beads with elution in 50 µl of TE (1X Tris-Ethylenediaminetetraacetic acid [EDTA], pH8). Capture hybridization of single or multiplexed up to 12 barcoded libraries (60 ng per sample) was performed using custom xGen^®^ Lockdown^®^ Probes (IDT; full details available at https://doi.org/10.35092/yhjc.11441001.v1). All 120-mer capture and blocking oligonucleotide probes were purchased from IDT as Ultramer DNA Oligos with Standard Desalting. Captured library fragments were further amplified by 12 cycles of aLM-PCR and run using an Agilent 2100 Bioanalyzer or TapeStation to estimate enrichment. Captured libraries were quantified by Qubit^®^ Fluorometer and quantitative Real Time-PCR (qRT-PCR) were used to dilute libraries for template preparation and Ion Sphere(tm) Particle (ISP) loading using Ion Chef(tm) System and sequencing on the Ion Proton(tm) platform with PI_v3_ semiconductor wafer chips per manufacturers recommended instructions. High-throughput sequencing of up to 39 multiplex captured libraries was carried out per PI_v3_ chip to achieve an average of 4.5 million reads per barcode. Reads containing the transposon IRDR element were processed using the SBCapSeq bioinformatic workflow as described(Mann et al., 2016b). For splink_454 sequencing (see **Supplementary Note 3**), SB insertion reads were generated by 454 GS Titanium sequencing (Roche) of pooled splinkerette PCR reactions with nested, barcoded primers was performed as described previously (Mann et al., 2012; Mann et al., 2015; March et al., 2011). Pre- and post-processing of 454 reads to assign sample DNA barcodes, filter out local hopping events from donor chromosomes, and map and orient the SB insertion sites across the entire nuclear genome of the mouse was performed. Donor chromosomes SB insertions were filtered away prior to identification of common insertion sites using SB Driver Analysis (Newberg et al., 2018a).

#### SB Driver Analysis Discovery, Progression and Trunk Drivers

Using the SB Driver Analysis framework (Newberg et al., 2018a), Discovery significant progression drivers are defined as statistically significant recurrently altered genes using False Discovery Rate (FDR; Benjamini–Hochberg procedure) multiple hypothesis testing *P*-value correction and Genome significant progression drivers are defined as statistically significant recurrently altered genes using Family Wise Error Rate (FWER; Holm–Bonferroni procedure) multiple hypothesis testing *P*-value correction from all SB insertion events within a dataset (no read depth filtering) to capture the widest possible list of potential driver genes. In contrast, Genome significant trunk drivers are defined as statistically significant recurrently altered genes using FWER multiple hypothesis testing *P*-value correction from SB insertion events with a pre-determined minimum read depth filtering of the dataset to report a statistically stringent prioritized driver gene list. Trunk drivers have the highest probability to be represented recurrently across tumors with the highest read depths, suggesting early and/or highly selected insertion events during bulk tumor growth. As a rule, Progression Drivers are a subset of Discovery Drivers and Trunk Drivers are often also Progression and/or Discovery Drivers (Newberg et al., 2018a). The BED files containing all SB insertions from histologically verified SB-Onc2.3 tumor specimens were filtered to consider only insertions represented by 100 or more reads in three or more tumors. When genes from a specimen genome were found to contain >1 SB insertion in the same RefSeq gene, only the SB insertion with the highest sequence read count was used for SB Driver Analysis calculations. SB Driver Analysis (Newberg et al., 2018a) was performed to identify the genes significantly enriched for insertions at high read counts with a FWER-corrected *P*<0.05 were termed trunk driver genes based on the inferred high clonality across the tumor population (Newberg et al., 2018a).

### qRT-PCR

Total RNA was purified and DNase treated using the RNeasy Mini Kit (Qiagen). Synthesis of cDNA was performed using SuperScript VILO Master Mix (Life Technologies). Quantitative PCR analysis was performed on the QuantStudio 12K Flex System (Life Technologies) or 7900HT Sequence Detection System (Applied Biosystem). All signals were normalized to the levels of GAPDH TaqMan probes. TaqMan probes were obtained from Life Technologies.

### Software

Unless otherwise noted, bioinformatic analysis pipelines, report generation, and figure visualization performed using bash, R, Python scripts, Vector NTI, and GraphPad Prism 8 software (Version 8.1.1).

### URLs

TgTn(sb-T2/Onc2.3) 14942Mbm, http://www.informatics.jax.org/allele/MGI:6441755; TgTn(sb-T2/Onc2.3)14922Mbm, http://www.informatics.jax.org/allele/MGI:6441757; TgTn(sb-T2/Onc2.3)14913Mbm, http://www.informatics.jax.org/allele/MGI:6441760; Mouse Genome Informatics (MGI) database, http://www.informatics.jax.org/; Gt(ROSA)26Sortm2(sb11)Njen, http://www.informatics.jax.org/allele/MGI:3839796; FVB/N-Tg(ACTB-cre)2Mrt/J, https://www.jax.org/strain/003376; FVB.129P2-Trp53tm1Brn, https://www.jax.org/strain/008462; 129S4-Trp53tm2Tyj, https://www.jax.org/strain/002101; Enrichr, http://amp.pharm.mssm.edu/Enrichr/; Tumor Suppressor Gene Database, https://bioinfo.uth.edu/TSGene/; UniProt, http://www.uniprot.org/; COSMIC Cancer Gene Census, https://cancer.sanger.ac.uk/census/; VENNY 2.1, https://bioinfogp.cnb.csic.es/tools/venny/; Oncoprinter, http://www.cbioportal.org/oncoprinter.

## Supplementary Notes

Supplementary Notes 1–3 may be downloaded from figshare http://dx.doi.org/10.6084/m9.figshare.12811979.

## Supplementary Tables

Supplementary Table datasets may be downloaded from figshare http://dx.doi.org/10.6084/m9.figshare.12811772. Brief descriptions of each table are provided for reference.

**Supplementary Table 1:** Tumor incidence and subgroup classifications by cohort groups

**Supplementary Table 2:** Sequencing projects summary by cohort

**Supplementary Table 3:** Specimen metafile data for projects sequenced using SBCapSeq protocol with Ion Torrent Proton sequencer

**Supplementary Table 4:** BED file of SB insertions for Skin-Onc2.3_SBC cohort

**Supplementary Table 5:** BED file of SB insertions for Lung-Onc2.3_SBC cohort

**Supplementary Table 6:** BED file of SB insertions for Lung-Onc3_SBC cohort

**Supplementary Table 7:** Specimen metafile data for projects sequenced using Splink_PCR protocol with 454 GS-FLX Titanium pyrosequencer

**Supplementary Table 8:** BED file of SB insertions for Lung29_454

**Supplementary Table 9:** SB Driver Analysis with discovery (FDR_hit = TRUE) and progression (FWER_hit = TRUE) for the Lung-Onc3_454 dataset (n=29)

**Supplementary Table 10:** BED file of SB insertions for HCA95_454

**Supplementary Table 11:** SB Driver Analysis with discovery (FDR_hit = TRUE) and progression (FWER_hit = TRUE) for the HAC-Onc3_454 dataset (n=95)

**Supplementary Table 12:** Trunk SB Driver Analysis with discovery (FDR_hit = TRUE) and progression (FWER_hit = TRUE) for the HAC-Onc3_454 dataset (n=95)

**Supplementary Table 13:** BED file of SB insertions for Liver-Onc2.3_SBC cohort

**Supplementary Table 14:** BED file of SB insertions for Other-Onc2.3_SBC cohort

**Supplementary Table 15:** BED file of SB insertions for Ovary-Onc2.3_SBC cohort

**Supplementary Table 16:** BED file of SB insertions for Onc2.3-Pan-Tumor-SBC27 cohort

**Supplementary Table 17:** Discovery and progression SB Driver Analysis for Onc2.3-Pan-Tumor-SBC27 cohort

**Supplementary Table 18:** Trunk SB Driver Analysis for Onc2.3-Pan-Tumor-SBC27. (Note: Six trunk drivers — *Aox2, Crlf3, Dgkd, Fgfr2, Ilf3*, and *Zfp292* — were manually removed from the dataset because supporting evidence from three related lung masses from a single animal provided insufficient evidence for tumor recurrence.)

**Supplementary Table 19:** Discovery and progression SB Driver Analysis for SB|Lung combined Onc2.3 and Onc3 cohorts

**Supplementary Table 20:** COSMIC v91 Cancer Gene Census TSG enrichment with SB-Onc2.3 Pan-tumor drivers

**Supplementary Table 21:** TSGdb enrichment with SB-Onc2.3 Pan-tumor drivers

**Supplementary Table 22:** Enrichr biological pathway and process analysis with SB-Onc2.3 Pan-tumor drivers

**Supplementary Table 23:** Literature References summarizing studies that provide in vivo evidence for validation of the TSGs identified

**Supplementary Table 24:** *Rasgrf1* gene expression analysis of from SB-Onc3 Lung tumor genomes

## Acknowledgments

The authors wish to thank the Copeland–Jenkins labs in Singapore and Houston, the K. Mann Lab at Moffitt Cancer Center and Martin McMahon and Aria Vaishnavi at Huntsman Cancer Institute for helpful discussions; we thank Gail Martin (UCSF) for Actin-beta-Cre mice; the Institute for Molecular and Cell Biology Histopathology Core; Adam Dupuy (Iowa) for providing the pT2/Onc2 plasmid; Holly Morris for husbandry and Debra Householder for genotype assistance of the T2/Onc2.3 transgenic founder mouse lines at NCI-Frederick (Frederick, MD, USA); Pearlyn Cheok, Nicole Lim, Dorothy Chen and Cherylin Wee for assistance with tumor monitoring and animal husbandry and Keith Rogers and Susan M. Rogers for assistance with necropsy and histotechnology services at IMCB (Singapore, Republic of Singapore); and Bethanie Gore for assistance with animal husbandry at MCC/USF (Tampa, FL, USA). Transgenic founder lines were created by the Mouse Cancer Genetics Program Transgenic Core Facility at NCI-Frederick. Necropsy and histology was performed by the Advanced Molecular Pathology Laboratory, Institute of Molecular and Cell Biology (A*STAR), Singapore and the Moffitt Tissue Core Facility. This work was supported by the Center for Cancer Research, National Cancer Institute (D.S., N.G.C., and N.A.J.), Biomedical Research Council, Agency for Science, Technology, and Research, Singapore (N.G.C. and N.A.J.), Cancer Prevention Research Institute of Texas (N.G.C. and N.A.J.), Cancer Center Support Grant (CCSG grant P30-CA76292) at the H. Lee Moffitt Cancer Center and Research Institute, a National Cancer Institute-designated Comprehensive Cancer Center (K.M.M. and M.B.M), and the Moffitt Skin Cancer SPORE (5P50CA168536-05) Career Enhancement Program Grant (M.B.M.).

## References

Abel, E.L., Angel, J.M., Kiguchi, K., and DiGiovanni, J. (2009). Multi-stage chemical carcinogenesis in mouse skin: fundamentals and applications. Nat Protoc 4, 1350–1362.

Aiderus, A., Newberg, J.Y., Guzman-Rojas, L., Contreras-Sandoval, A.M., Meshey, A.L., Jones, D.J., Amaya-Manzanares, F., Rangel, R., Ward, J.M., Lee, S.-C., et al. (2019). Transposon mutagenesis identifies cooperating genetic drivers during keratinocyte transformation and cutaneous squamous cell carcinoma progression. bioRxiv, 2019.2012.2024.887968.

Andricovich, J., Perkail, S., Kai, Y., Casasanta, N., Peng, W., and Tzatsos, A. (2018). Loss of KDM6A Activates Super-Enhancers to Induce Gender-Specific Squamous-like Pancreatic Cancer and Confers Sensitivity to BET Inhibitors. Cancer Cell 33, 512–526 e518.

Bamford, S., Dawson, E., Forbes, S., Clements, J., Pettett, R., Dogan, A., Flanagan, A., Teague, J., Futreal, P.A., Stratton, M.R., et al. (2004). The COSMIC (Catalogue of Somatic Mutations in Cancer) database and website. Br J Cancer 91, 355–358.

Bard-Chapeau, E.A., Nguyen, A.T., Rust, A.G., Sayadi, A., Lee, P., Chua, B.Q., New, L.S., de Jong, J., Ward, J.M., Chin, C.K., et al. (2014). Transposon mutagenesis identifies genes driving hepatocellular carcinoma in a chronic hepatitis B mouse model. Nat Genet 46, 24–32.

Chen, E.Y., Tan, C.M., Kou, Y., Duan, Q., Wang, Z., Meirelles, G.V., Clark, N.R., and Ma’ayan, A. (2013). Enrichr: interactive and collaborative HTML5 gene list enrichment analysis tool. BMC Bioinformatics 14, 128.

Chitsazzadeh, V., Coarfa, C., Drummond, J.A., Nguyen, T., Joseph, A., Chilukuri, S., Charpiot, E., Adelmann, C.H., Ching, G., Nguyen, T.N., et al. (2016). Cross-species identification of genomic drivers of squamous cell carcinoma development across preneoplastic intermediates. Nat Commun 7, 12601.

Cho, S.J., Yoon, C., Lee, J.H., Chang, K.K., Lin, J.X., Kim, Y.H., Kook, M.C., Aksoy, B.A., Park, D.J., Ashktorab, H., et al. (2018). KMT2C Mutations in Diffuse-Type Gastric Adenocarcinoma Promote Epithelial-to-Mesenchymal Transition. Clin Cancer Res 24, 6556–6569.

Collier, L.S., Carlson, C.M., Ravimohan, S., Dupuy, A.J., and Largaespada, D.A. (2005). Cancer gene discovery in solid tumours using transposon-based somatic mutagenesis in the mouse. Nature 436, 272–276.

Collier, L.S., and Largaespada, D.A. (2007). Transposons for cancer gene discovery: Sleeping Beauty and beyond. Genome Biol 8 Suppl 1, S15.

Copeland, N.G., and Jenkins, N.A. (2010). Harnessing transposons for cancer gene discovery. Nat Rev Cancer 10, 696–706.

Dhar, S.S., Zhao, D., Lin, T., Gu, B., Pal, K., Wu, S.J., Alam, H., Lv, J., Yun, K., Gopalakrishnan, V., et al. (2018). MLL4 Is Required to Maintain Broad H3K4me3 Peaks and Super-Enhancers at Tumor Suppressor Genes. Mol Cell 70, 825–841 e826.

Dorr, C., Janik, C., Weg, M., Been, R.A., Bader, J., Kang, R., Ng, B., Foran, L., Landman, S.R., O’Sullivan, M.G., et al. (2015). Transposon Mutagenesis Screen Identifies Potential Lung Cancer Drivers and CUL3 as a Tumor Suppressor. Mol Cancer Res 13, 1238–1247.

Dupuy, A.J., Akagi, K., Largaespada, D.A., Copeland, N.G., and Jenkins, N.A. (2005). Mammalian mutagenesis using a highly mobile somatic Sleeping Beauty transposon system. Nature 436, 221–226.

Dupuy, A.J., Rogers, L.M., Kim, J., Nannapaneni, K., Starr, T.K., Liu, P., Largaespada, D.A., Scheetz, T.E., Jenkins, N.A., and Copeland, N.G. (2009). A modified sleeping beauty transposon system that can be used to model a wide variety of human cancers in mice. Cancer Res 69, 8150–8156.

Forbes, S.A., Tang, G., Bindal, N., Bamford, S., Dawson, E., Cole, C., Kok, C.Y., Jia, M., Ewing, R., Menzies, A., et al. (2010). COSMIC (the Catalogue of Somatic Mutations in Cancer): a resource to investigate acquired mutations in human cancer. Nucleic Acids Res 38, D652–657.

Haigis, K.M. (2017). KRAS Alleles: The Devil Is in the Detail. Trends Cancer 3, 686–697.

Hayashi, T., Desmeules, P., Smith, R.S., Drilon, A., Somwar, R., and Ladanyi, M. (2018). RASA1 and NF1 are Preferentially Co-Mutated and Define A Distinct Genetic Subset of Smoking-Associated Non-Small Cell Lung Carcinomas Sensitive to MEK Inhibition. Clin Cancer Res 24, 1436–1447.

Higa, K.C., and DeGregori, J. (2019). Decoy fitness peaks, tumor suppression, and aging. Aging Cell 18, e12938.

Huang, J., Chen, M., Xu, E.S., Luo, L., Ma, Y., Huang, W., Floyd, W., Klann, T.S., Kim, S.Y., Gersbach, C.A., et al. (2019). Genome-wide CRISPR Screen to Identify Genes that Suppress Transformation in the Presence of Endogenous Kras(G12D). Sci Rep 9, 17220.

Ichise, T., Yoshida, N., and Ichise, H. (2019). CBP/p300 antagonises EGFR-Ras-Erk signalling and suppresses increased Ras-Erk signalling-induced tumour formation in mice. J Pathol 249, 39–51.

Ivics, Z., Hackett, P.B., Plasterk, R.H., and Izsvak, Z. (1997). Molecular reconstruction of Sleeping Beauty, a Tc1-like transposon from fish, and its transposition in human cells. Cell 91, 501–510.

Izsvak, Z., and Ivics, Z. (2004). Sleeping beauty transposition: biology and applications for molecular therapy. Mol Ther 9, 147–156.

Izsvak, Z., Ivics, Z., and Plasterk, R.H. (2000). Sleeping Beauty, a wide host-range transposon vector for genetic transformation in vertebrates. J Mol Biol 302, 93–102.

Izumchenko, E., Chang, X., Brait, M., Fertig, E., Kagohara, L.T., Bedi, A., Marchionni, L., Agrawal, N., Ravi, R., Jones, S., et al. (2015). Targeted sequencing reveals clonal genetic changes in the progression of early lung neoplasms and paired circulating DNA. Nat Commun 6, 8258.

Jia, D., Augert, A., Kim, D.W., Eastwood, E., Wu, N., Ibrahim, A.H., Kim, K.B., Dunn, C.T., Pillai, S.P.S., Gazdar, A.F., et al. (2018). Crebbp Loss Drives Small Cell Lung Cancer and Increases Sensitivity to HDAC Inhibition. Cancer Discov 8, 1422–1437.

Jonkers, J., Meuwissen, R., van der Gulden, H., Peterse, H., van der Valk, M., and Berns, A. (2001). Synergistic tumor suppressor activity of BRCA2 and p53 in a conditional mouse model for breast cancer. Nat Genet 29, 418–425.

Kas, S.M., de Ruiter, J.R., Schipper, K., Schut, E., Bombardelli, L., Wientjens, E., Drenth, A.P., de Korte-Grimmerink, R., Mahakena, S., Phillips, C., et al. (2018). Transcriptomics and Transposon Mutagenesis Identify Multiple Mechanisms of Resistance to the FGFR Inhibitor AZD4547. Cancer Res 78, 5668–5679.

Katigbak, A., Cencic, R., Robert, F., Senecha, P., Scuoppo, C., and Pelletier, J. (2016). A CRISPR/Cas9 Functional Screen Identifies Rare Tumor Suppressors. Sci Rep 6, 38968.

Keng, V.W., Sia, D., Sarver, A.L., Tschida, B.R., Fan, D., Alsinet, C., Sole, M., Lee, W.L., Kuka, T.P., Moriarity, B.S., et al. (2013). Sex bias occurrence of hepatocellular carcinoma in Poly7 molecular subclass is associated with EGFR. Hepatology 57, 120–130.

Kodama, T., Yi, J., Newberg, J.Y., Tien, J.C., Wu, H., Finegold, M.J., Kodama, M., Wei, Z., Tamura, T., Takehara, T., et al. (2018). Molecular profiling of nonalcoholic fatty liver disease-associated hepatocellular carcinoma using SB transposon mutagenesis. Proc Natl Acad Sci U S A 115, E10417–E10426.

Kuleshov, M.V., Jones, M.R., Rouillard, A.D., Fernandez, N.F., Duan, Q., Wang, Z., Koplev, S., Jenkins, S.L., Jagodnik, K.M., Lachmann, A., et al. (2016). Enrichr: a comprehensive gene set enrichment analysis web server 2016 update. Nucleic Acids Res 44, W90–97.

Larrieu, D., Brunet, M., Vargas, C., Hanoun, N., Ligat, L., Dagnon, L., Lulka, H., Pommier, R.M., Selves, J., Jady, B.E., et al. (2020). The E3 ubiquitin ligase TRIP12 participates in cell cycle progression and chromosome stability. Sci Rep 10, 789.

Lewandoski, M., Meyers, E.N., and Martin, G.R. (1997). Analysis of Fgf8 gene function in vertebrate development. Cold Spring Harb Symp Quant Biol 62, 159–168.

Liang, Y.N., Liu, Y., Meng, Q., Li, X., Wang, F., Yao, G., Wang, L., Fu, S., and Tong, D. (2015). RBMS3 is a tumor suppressor gene that acts as a favorable prognostic marker in lung squamous cell carcinoma. Med Oncol 32, 459.

Mann, K.M., Jenkins, N.A., Copeland, N.G., and Mann, M.B. (2014a). Transposon insertional mutagenesis models of cancer. Cold Spring Harb Protoc 2014, 235–247.

Mann, K.M., Mann, M.B., Guzman-Rojas, L., Amaya-Manzanares, F., Jones, D.J., Newberg, J.Y., Jenkins, N.A., and Copeland, N.G. (2016a). SBCapSeq Protocol: a method for selective cloning of Sleeping Beauty transposon insertions using liquid capture hybridization and Ion Torrent semiconductor sequencing. Protocol Exchange.

Mann, K.M., Newberg, J.Y., Black, M.A., Jones, D.J., Amaya-Manzanares, F., Guzman-Rojas, L., Kodama, T., Ward, J.M., Rust, A.G., van der Weyden, L., et al. (2016b). Analyzing tumor heterogeneity and driver genes in single myeloid leukemia cells with SBCapSeq. Nat Biotechnol 34, 962–972.

Mann, K.M., Ward, J.M., Yew, C.C., Kovochich, A., Dawson, D.W., Black, M.A., Brett, B.T., Sheetz, T.E., Dupuy, A.J., Australian Pancreatic Cancer Genome, I., et al. (2012). Sleeping Beauty mutagenesis reveals cooperating mutations and pathways in pancreatic adenocarcinoma. Proc Natl Acad Sci U S A 109, 5934–5941.

Mann, M.B., Black, M.A., Jones, D.J., Ward, J.M., Yew, C.C., Newberg, J.Y., Dupuy, A.J., Rust, A.G., Bosenberg, M.W., McMahon, M., et al. (2015). Transposon mutagenesis identifies genetic drivers of Braf(V600E) melanoma. Nat Genet 47, 486–495.

Mann, M.B., Jenkins, N.A., Copeland, N.G., and Mann, K.M. (2014b). Sleeping Beauty mutagenesis: exploiting forward genetic screens for cancer gene discovery. Curr Opin Genet Dev 24, 16–22.

Mann, M.B., Mann, K.M., Contreras-Sandoval, A.M., Guzman-Rojas, L., Newberg, J.Y., Meshey, A.L., Jones, D.J., Amaya-Manzanares, F., Copeland, N.G., and Jenkins, N.A. (2019). SBCapSeq Protocol manuscript files for ‘Quantifying tumor heterogeneity, clonal dynamics, and cancer driver gene evolution from Sleeping Beauty transposon mutagenesis models using SBCapSeq’. National Institutes of Health Dataset (NIHFigsharecom).

March, H.N., Rust, A.G., Wright, N.A., ten Hoeve, J., de Ridder, J., Eldridge, M., van der Weyden, L., Berns, A., Gadiot, J., Uren, A., et al. (2011). Insertional mutagenesis identifies multiple networks of cooperating genes driving intestinal tumorigenesis. Nature genetics 43, 1202–1209.

Martincorena, I., Fowler, J.C., Wabik, A., Lawson, A.R.J., Abascal, F., Hall, M.W.J., Cagan, A., Murai, K., Mahbubani, K., Stratton, M.R., et al. (2018). Somatic mutant clones colonize the human esophagus with age. Science 362, 911–917.

Martincorena, I., Roshan, A., Gerstung, M., Ellis, P., Van Loo, P., McLaren, S., Wedge, D.C., Fullam, A., Alexandrov, L.B., Tubio, J.M., et al. (2015). Tumor evolution. High burden and pervasive positive selection of somatic mutations in normal human skin. Science 348, 880–886.

Mathur, R., Alver, B.H., San Roman, A.K., Wilson, B.G., Wang, X., Agoston, A.T., Park, P.J., Shivdasani, R.A., and Roberts, C.W. (2017). ARID1A loss impairs enhancer-mediated gene regulation and drives colon cancer in mice. Nat Genet 49, 296–302.

Molina-Sanchez, P., and Lujambio, A. (2019). Experimental Models for Preclinical Research in Hepatocellular Carcinoma. In Hepatocellular Carcinoma: Translational Precision Medicine Approaches, Y. Hoshida, ed. (Cham (CH)), pp. 333–358.

Newberg, J.Y., Black, M.A., Jenkins, N.A., Copeland, N.G., Mann, K.M., and Mann, M.B. (2018a). SB Driver Analysis: a Sleeping Beauty cancer driver analysis framework for identifying and prioritizing experimentally actionable oncogenes and tumor suppressors. Nucleic Acids Res 46, e94.

Newberg, J.Y., Mann, K.M., Mann, M.B., Jenkins, N.A., and Copeland, N.G. (2018b). SBCDDB: Sleeping Beauty Cancer Driver Database for gene discovery in mouse models of human cancers. Nucleic Acids Res 46, D1011–D1017.

O’Donnell, K.A., Keng, V.W., York, B., Reineke, E.L., Seo, D., Fan, D., Silverstein, K.A., Schrum, C.T., Xie, W.R., Mularoni, L., et al. (2012). A Sleeping Beauty mutagenesis screen reveals a tumor suppressor role for Ncoa2/Src-2 in liver cancer. Proc Natl Acad Sci U S A 109, E1377–1386.

Olive, K.P., Tuveson, D.A., Ruhe, Z.C., Yin, B., Willis, N.A., Bronson, R.T., Crowley, D., and Jacks, T. (2004). Mutant p53 gain of function in two mouse models of Li-Fraumeni syndrome. Cell 119, 847–860.

Perez-Mancera, P.A., Rust, A.G., van der Weyden, L., Kristiansen, G., Li, A., Sarver, A.L., Silverstein, K.A., Grutzmann, R., Aust, D., Rummele, P., et al. (2012). The deubiquitinase USP9X suppresses pancreatic ductal adenocarcinoma. Nature 486, 266–270.

Perna, D., Karreth, F.A., Rust, A.G., Perez-Mancera, P.A., Rashid, M., Iorio, F., Alifrangis, C., Arends, M.J., Bosenberg, M.W., Bollag, G., et al. (2015). BRAF inhibitor resistance mediated by the AKT pathway in an oncogenic BRAF mouse melanoma model. Proc Natl Acad Sci U S A 112, E536–545.

Rangel, R., Lee, S.C., Hon-Kim Ban, K., Guzman-Rojas, L., Mann, M.B., Newberg, J.Y., Kodama, T., McNoe, L.A., Selvanesan, L., Ward, J.M., et al. (2016). Transposon mutagenesis identifies genes that cooperate with mutant Pten in breast cancer progression. Proc Natl Acad Sci U S A 113, E7749–E7758.

Rogers, L.M., Olivier, A.K., Meyerholz, D.K., and Dupuy, A.J. (2013). Adaptive immunity does not strongly suppress spontaneous tumors in a Sleeping Beauty model of cancer. J Immunol 190, 4393–4399.

Senft, D., Qi, J., and Ronai, Z.A. (2018). Ubiquitin ligases in oncogenic transformation and cancer therapy. Nat Rev Cancer 18, 69–88.

Sondka, Z., Bamford, S., Cole, C.G., Ward, S.A., Dunham, I., and Forbes, S.A. (2018). The COSMIC Cancer Gene Census: describing genetic dysfunction across all human cancers. Nat Rev Cancer 18, 696–705.

Starr, T.K., Allaei, R., Silverstein, K.A., Staggs, R.A., Sarver, A.L., Bergemann, T.L., Gupta, M., O’Sullivan, M.G., Matise, I., Dupuy, A.J., et al. (2009). A transposon-based genetic screen in mice identifies genes altered in colorectal cancer. Science 323, 1747–1750.

Suh, J.H., Johnson, A., Albacker, L., Wang, K., Chmielecki, J., Frampton, G., Gay, L., Elvin, J.A., Vergilio, J.A., Ali, S., et al. (2016). Comprehensive Genomic Profiling Facilitates Implementation of the National Comprehensive Cancer Network Guidelines for Lung Cancer Biomarker Testing and Identifies Patients Who May Benefit From Enrollment in Mechanism-Driven Clinical Trials. Oncologist 21, 684–691.

Takeda, H., Wei, Z., Koso, H., Rust, A.G., Yew, C.C., Mann, M.B., Ward, J.M., Adams, D.J., Copeland, N.G., and Jenkins, N.A. (2015). Transposon mutagenesis identifies genes and evolutionary forces driving gastrointestinal tract tumor progression. Nat Genet 47, 142–150.

Westcott, P.M., Halliwill, K.D., To, M.D., Rashid, M., Rust, A.G., Keane, T.M., Delrosario, R., Jen, K.Y., Gurley, K.E., Kemp, C.J., et al. (2015). The mutational landscapes of genetic and chemical models of Kras-driven lung cancer. Nature 517, 489–492.

Wu, Q., Tian, Y., Zhang, J., Tong, X., Huang, H., Li, S., Zhao, H., Tang, Y., Yuan, C., Wang, K., et al. (2018). In vivo CRISPR screening unveils histone demethylase UTX as an important epigenetic regulator in lung tumorigenesis. Proc Natl Acad Sci U S A 115, E3978–E3986.

Yokoyama, A., Kakiuchi, N., Yoshizato, T., Nannya, Y., Suzuki, H., Takeuchi, Y., Shiozawa, Y., Sato, Y., Aoki, K., Kim, S.K., et al. (2019). Age-related remodelling of oesophageal epithelia by mutated cancer drivers. Nature 565, 312–317.

Zender, L., Xue, W., Zuber, J., Semighini, C.P., Krasnitz, A., Ma, B., Zender, P., Kubicka, S., Luk, J.M., Schirmacher, P., et al. (2008). An oncogenomics-based in vivo RNAi screen identifies tumor suppressors in liver cancer. Cell 135, 852–864.

Zhang, D.D., Lo, S.C., Sun, Z., Habib, G.M., Lieberman, M.W., and Hannink, M. (2005). Ubiquitination of Keap1, a BTB-Kelch substrate adaptor protein for Cul3, targets Keap1 for degradation by a proteasome-independent pathway. J Biol Chem 280, 30091–30099.

Zhao, M., Kim, P., Mitra, R., Zhao, J., and Zhao, Z. (2016). TSGene 2.0: an updated literature-based knowledgebase for tumor suppressor genes. Nucleic Acids Res 44, D1023–1031.

Zhao, M., Sun, J., and Zhao, Z. (2013). TSGene: a web resource for tumor suppressor genes. Nucleic Acids Res 41, D970–976.

